# Liver-specific loss of the transcriptional coregulator ARGLU1 protects against diet-induced obesity in mice through decreased lipid absorption

**DOI:** 10.64898/2026.02.03.703579

**Authors:** Sarah B. Cash, Michael F. Saikali, Lilia Magomedova, Sarah A. Shawky, Fatemeh Molaei, Jin Shi, Pierre E. Thibeault, Sulayman A. Lyons, Candice Kwok, Taira Kiyota, Stéphane Angers, Ahmed Aman, Jacqueline L. Beaudry, Carolyn L. Cummins

## Abstract

New medications that decrease food intake show promise in addressing the global obesity epidemic; however, additional therapies are needed for patients who are intolerant or unresponsive to these drugs. ARGLU1 is a transcriptional coregulator of several nuclear receptors. Herein, we detail the discovery that mice lacking ARGLU1 in hepatocytes (LKO) are resistant to diet-induced obesity, with no difference in food intake, locomotion, or energy expenditure compared to wildtype mice. Interestingly, LKO mice exhibited decreased CYP8B1, corresponding to fewer 12α-hydroxylated bile acids, increasing the relative hydrophobicity of bile which decreases lipid emulsification. Notably, LKO mice have lower plasma and liver cholesterol levels, attributed to impaired lipid absorption. Furthermore, LKO mice showed preferential use of fatty acids as their fuel source. Herein, we establish a mechanistic basis for resistance of ARGLU1 LKO mice to diet-induced obesity, identifying a potential new molecular target for obesity.

## INTRODUCTION

Obesity is a growing global health concern^1,2^ that can result from dysfunctional intermediary metabolism, affecting the regulation of fatty acids, cholesterol, and glucose.^3^ Although current pharmacological treatments for obesity exist, the growing incidence rates necessitate the development of additional therapeutic options.^4^

Bile acids are amphipathic molecules critical for the emulsification of dietary lipids required for intestinal lipid absorption. Transcription factors can directly and indirectly regulate bile acid homeostasis (i.e., bile acid synthesis, hepatic uptake and efflux, and intestinal reabsorption.^5-11^ In the liver, bile acids are synthesized from cholesterol in either the classical pathway to generate cholic acid (CA) or from the alternative pathway to generate chenodeoxycholic acid (CDCA). In rodents, CDCA can be further modified to form muricholic acid (MCA). The majority of the bile acid pool in mice is taurine-conjugated, primarily consisting of Tauro-CA (TCA) and Tauro-αβωMCA (TαβωMCA).^12^ In the intestine, endogenously produced primary bile acids can be converted into secondary bile acids by intestinal microbes.

The enzyme CYP7A1 catalyzes the first rate-limiting step of bile acid synthesis whereas CYP8B1 is essential for 12α-hydroxylation of bile acids. Thus, CYP8B1 dictates the ratio of 12α-hydroxylated to non-12α-hydroxylated bile acids.^13-15^ As bile acids possess both hydrophobic and hydrophilic characteristics, it is the orientation of the hydroxyl group (i.e., on either the α-face or β-face) that dictates the overall hydrophobicity. For example, the hydrophobic 12α-hydroxylated bile acid TCA is more efficient at lipid emulsification compared to the non-12α-hydroxylated bile acids TαβωMCA.^14^ *Cyp8b1^-/-^* mice have decreased hydrophobic 12α-hydroxylated bile acids, resulting in more hydrophilic bile composition (i.e., TαβωMCA) that is less efficient at absorbing dietary lipids.^13,15^ Thus, changes to bile acid pool composition have been considered as a mechanism to influence lipid absorption and reduce diet-induced obesity (DIO).^13,15-17^

Arginine- and glutamate-rich 1 (ARGLU1) is a transcriptional coregulator for the glucocorticoid receptor and is essential for alternative splicing within neuronal tissues.^18^ ARGLU1 has also been shown to directly interact with Mediator subunit 1 (MED1) in the context of estrogen receptor activation.^19^ Given ARGLU1’s role as a coregulator for several metabolically important nuclear receptors (NRs) *in vitro*^18^ and widespread tissue expression, including the liver, we investigated whether hepatic ARGLU1 plays a role in regulating lipid metabolism and bile acid homeostasis. Herein, we demonstrate that ARGLU1 is crucial for the synthesis of 12α-hydroxylated bile acids, therefore impacting the efficiency of dietary lipid absorption.

## RESULTS

### Male and female ARGLU1 LKO mice are resistant to DIO, independent of food intake, locomotion, and energy expenditure

To generate a DIO model, we placed female and male WT and ARGLU1 LKO mice on a high-fat, high-cholesterol diet (HFD, 4.5 kcal/g; **Table S1**) for 12-weeks and compared them to mice fed a chow diet (3 kcal/g) (**Figure 1A**). As expected, male and female WT mice demonstrate a significant increase in body weight compared to chow diet (**Figure 1B-C**, **1J-K**). Notably, in both sexes, LKO mice weighed less (females: 13%, males: 20%) than WT mice at 12- weeks, demonstrating resistance to DIO (**Figure 1B-C**, **1J-K**). On chow diet, there was no difference in body weight between WT and LKO mice, indicating that LKO alters weight gain only in obese mice (**Figure 1B-C**, **1J-K**). We next measured fat distribution in LKO mice versus WT mice, examining adipose tissue weight and body fat composition using dual-energy X-ray absorptiometry (DEXA). DEXA analysis revealed that female HFD-fed LKO mice had decreased visceral and total fat area compared to HFD-fed WT mice (**Figure 1D**); however, male LKO mice had similar visceral fat area compared to HFD-fed WT mice (**Figure 1L**). In male WT mice, total fat area was increased 48% upon HFD-feeding (*P*<0.05), but this increase was not significant in male LKO mice upon HFD-feeding (**Figure 1L**). Consistent with decreased body weight, both female and male LKO mice exhibited reduced adipose depot weights compared to HFD-fed WT mice (**Figure 1E-H, 1M-P**). Liver weight was significantly increased in HFD-fed WT mice in both sexes; however, this was not observed in LKO mice upon HFD feeding (**Figure 1I**, **1Q**). Changes in organ weights were not due to overall differences in growth as assessed by femur length (male: 15.9 ± 0.1 mm vs 16.0 ± 0.1 mm; female: 15.4 ± 0.2 mm vs 15.2 ± 0.4 mm, for WT HFD and LKO HFD, respectively). We determined that the difference in body weight gain was not due to the Albumin-Cre promoter by comparing body weight gain between Albumin^Cre/+^ and Arglu1^fl/fl^ mice (**Table S2**).

**Figure 1.**
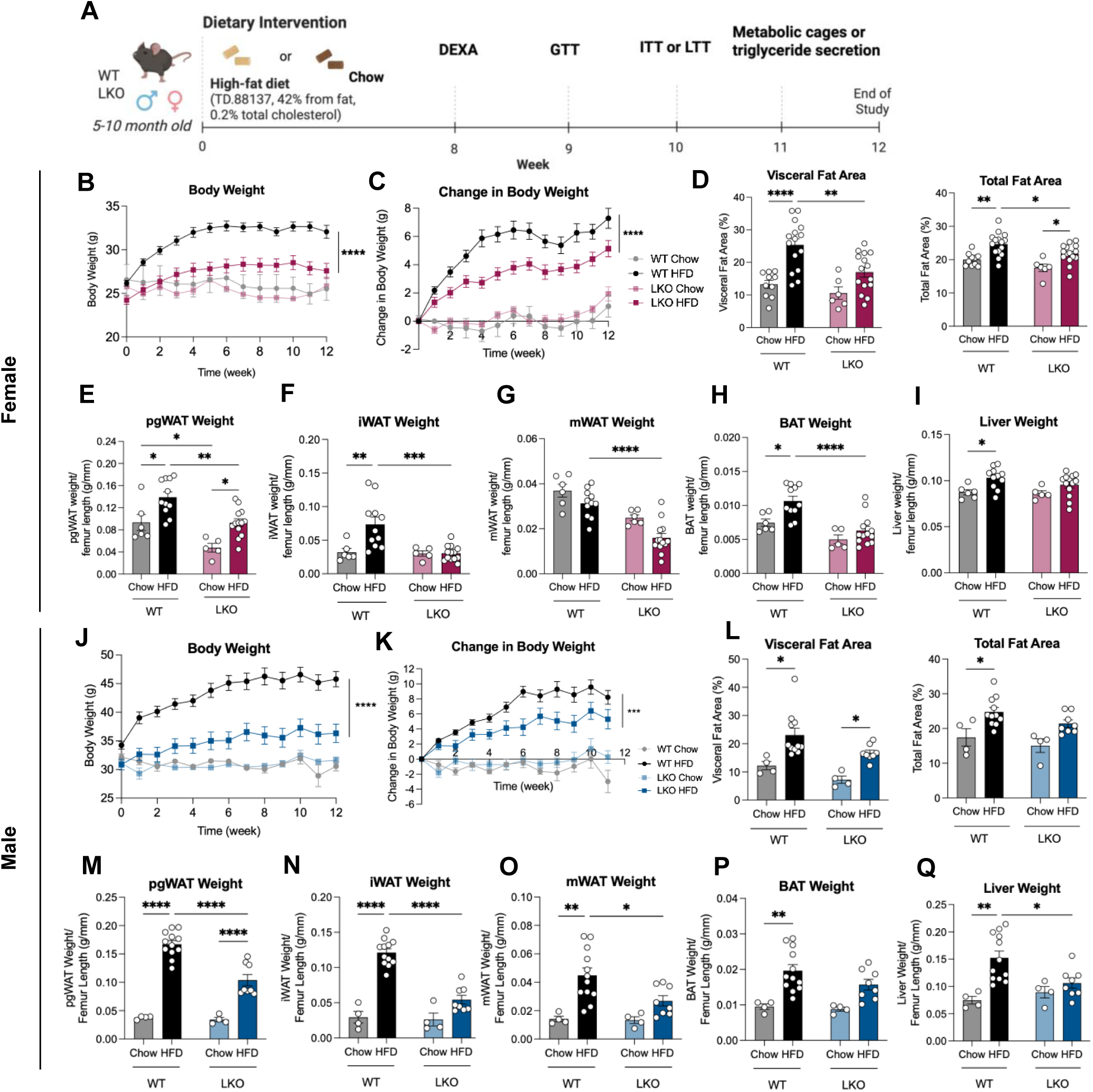
ARGLU1 LKO mice are resistant to diet-induced obesity. (A) Schematic of experiments performed during 12 weeks of HFD. Body weight (B, J) and changes in body weight (C, K) were measured from female and male mice. Visceral and total fat area was measured in female (C) and male (L) mice using dual-energy X-ray absorptiometry. Upon sacrifice, tissue weights from perigonadal WAT (pgWAT) (E, M), inguinal WAT (iWAT) (F, N), mesenteric (mWAT) (G, O), and brown adipose tissue (BAT) (H, P) were measured in female and male mice and normalized to femur length from each mouse. Liver weight was also compared to femur length in female (I) and male (Q) mice. \**P* ≤ 0.05, \*\**P* ≤ 0.01, \*\*\**P* ≤ 0.001, \*\*\*\**P* ≤ 0.0001, by two-way ANOVA followed by Holm-Sidak test (B-Q). Data are represented as mean ± SEM with individual animals noted as dots (N=4-17).

We next performed glucose and insulin tolerance tests, dosing based on lean body mass. Glucose (**Figure S1A**) and insulin (**Figure S1B**) tolerance did not differ between female HFD-fed WT and LKO mice; however, male HFD-fed LKO mice showed significantly improved glucose tolerance compared to WT mice (**Figure S1C**). There was no difference in insulin tolerance between WT mice and LKO mice upon HFD-feeding (**Figure S1D**). Furthermore, male LKO mice had lower fed (decreased 48%, *P*<0.05; **Figure S1E**) and fasted (decreased 66%, *P*<0.05; **Figure S1F**) plasma insulin levels compared to WT mice.

To determine the mechanism of the resistance to DIO in LKO mice, we monitored food intake. In both sexes, WT and LKO mice consumed similar daily and cumulative calories from the diet, indicating that the weight differences were not attributed to food intake (**Figure S2A-B**, **S2E-F**). Next, we examined locomotion and found no differences in total or pedestrian locomotion between WT and LKO mice (**Figure S2C**, **S2G**). To assess whether differences in total and basal energy were contributing to changes in body weight, we housed the mice at thermoneutrality and placed them in metabolic cages. No differences in energy expenditure were found between WT and LKO HFD-fed mice in either sex after accounting for body weight as a variable using ANCOVA (**Figure S2D**, **S2H**). These findings suggest that the resistance to DIO in LKO mice is not due to differences in food intake, locomotion, or energy expenditure.

### LKO HFD-fed mice excrete more fecal lipids and have improved lipid tolerance, compared to HFD-fed WT mice

To reconcile the lack of differences in food intake, locomotion, or energy expenditure with the changes observed in body weight, we considered whether there were differences in lipid absorption. While both sexes of WT and LKO mice had similar lipid intake (**Figure 2A**, **2I**), LKO mice had increased fecal lipid output compared to WT mice (**Figure 2B**, **2J**), corresponding to decreased dietary lipid absorption (**Figure 2C**, **2K**). Further analysis revealed increased cholesterol and triglyceride content in feces of both female (**Figure 2D-E**) and male (**Figure 2L-M**) LKO mice compared to WT controls. To further evaluate this finding, male and female HFD-fed WT and LKO mice underwent a lipid tolerance test (LTT) (**Figure 2F, 2N**). Male LKO mice showed reduced peak plasma triglyceride levels compared to WT mice, resulting in significantly lower AUCs compared to WT (**Figure 2N**). In contrast, no significant differences were detected in female mice between genotypes in peak plasma or area under the curve (AUC) (**Figure 2F**). Literature revealed that female mice have compromised sensitivity to the LTT due to increased basal systemic lipoprotein lipase (LPL) levels compared to male mice.^20^ This is consistent with our findings that demonstrate peak plasma triglyceride levels of 90 mg/dL in WT female mice compared to 350 mg/dL in WT male mice during LTT. Intestinal length did not differ between genotypes (data not shown).

**Figure 2.**
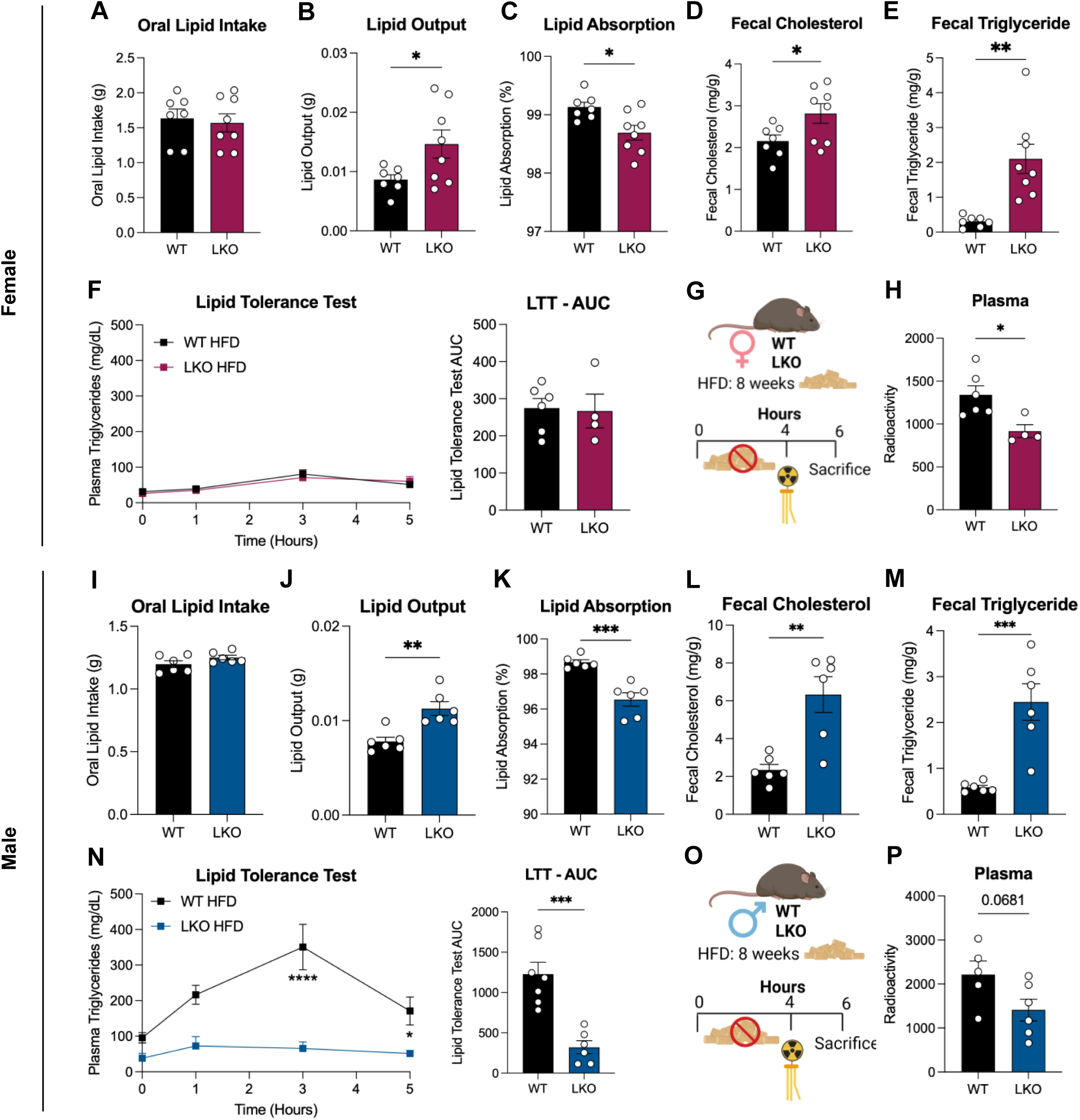
HFD-fed LKO mice have decreased lipid absorption compared to WT mice. Oral lipid intake was measured in singly housed female (A) and male (I) mice (N=6-8) for a 72-hour period, in the final week of the study. Using the Folch lipid extraction method on stool samples, lipid output (B, J) and corresponding lipid absorption (C, K) was measured in HFD-fed female and male fecal samples (N=6-8). Both fecal cholesterol (D, L) and triglyceride (E, M) were measured in female and male mice (N=5-8). Overnight fasted female (F) and male (N) mice (N=4-7) were dosed with 200 µL of olive oil and plasma triglyceride content was measured over a 5-hour-period, area under the curve is reported. Fasted (4-hour) mice (N=4-6) were dosed with 200 µL of olive oil with 1 µCi ^14^C-Triolein (G, O) in female (H) and male (P) mice, plasma was collected 2 hours later. LTT, lipid tolerance test; CPM, counts per minute. \**P* ≤ 0.05, \*\**P* ≤ 0.01, \*\*\**P* ≤ 0.001, \*\*\*\**P* ≤ 0.0001, by two-way ANOVA followed by Holm-Sidak test (F, N) or by unpaired two-tailed T-Test (A-E, G-M, O-P). Data are represented as mean ± SEM with individual animals noted as dots.

To directly assess lipid absorption, we dosed 4-hour fasted mice with ^14^C-triolein and measured plasma radioactivity after 2 hours (**Figure 2G**, **2O**). Female LKO mice had significantly decreased plasma ^14^C activity compared to WT mice (**Figure 2H**). In male mice, plasma radioactivity was not significantly different between WT and LKO mice but tended to be lower in LKO mice (*P* = 0.068; **Figure 2P**). We measured gene expression of key chylomicron formation or cholesterol and fatty acid transporters in the duodenum and jejunum and found no decrease in transporter expression in LKO mice versus WT mice that would be responsible for reduced fatty acid or cholesterol uptake (**Figure S3A-F**). Thus, we conclude LKO mice have reduced lipid absorption, though the mechanism needed to be further investigated.

### Transcriptome analysis revealed altered bile acid and cholesterol pathways in LKO mice, compared to WT mice

To further investigate the mechanism underlying DIO resistance in LKO mice, we performed transcriptomic analysis on female liver samples from chow and HFD-fed WT and LKO mice. Principal component analysis (PCA) revealed distinct genotype clustering (**Figure 3A**), with transcriptomic responses to HFD feeding differing between genotypes, with minimal overlap in shared HFD-induced differentially expressed genes (**Figure S4A**). We identified 527 downregulated and 1226 upregulated genes in HFD-fed LKO mice compared to HFD-fed WT controls (**Figure 3B**, adjusted *P-*value ≤0.05 and log_2_ fold change ≥|1.5|). Upregulated gene ontology pathways in HFD-fed LKO mice compared to WT mice included fatty acid and carboxylic acid metabolic processes (**Figure 3C, Table S3**), while downregulated pathways included steroid metabolic processes (**Figure 3D, Table S3**). In chow-fed LKO mice, we found 1192 genes significantly upregulated, and 629 genes downregulated, compared to chow-fed WT mice (**Figure S4B**). In chow-fed LKO mice, fatty acid metabolism pathways were upregulated, while cholesterol metabolism and bile acid biosynthetic pathways were downregulated compared to chow-fed WT mice (**Figure S4C**-**D**). KEGG pathway analysis revealed significant downregulation of bile secretion and steroid hormone biosynthesis pathways in LKO mice compared to WT mice in both HFD (**Figure 3E**) and chow-fed conditions (**Figure S4E**). From our transcriptomic profile of the WT and LKO HFD-fed samples, C/EBP emerged as the highest scoring transcription factor of interest due to its enriched motifs in promoter regions of significantly downregulated genes that were identified in LKO mice compared to WT mice (**Figure 3F**). Gene-set enrichment analysis showed a significant reduction in the bile acid metabolism pathway in HFD-fed LKO mice compared to WT mice (**Figure 3G**). Analysis of differentially expressed genes revealed several trends in LKO mice, including increased fatty acid oxidation, decreased bile acid and cholesterol hepatic transporters, and decreased bile acid synthesis (**Figure 3H**). Bile acid sulfonation enzymes were significantly decreased in LKO mice, while bile acid glucuronidation enzymes were significantly increased compared to WT mice (**Figure 3H**). Collectively, these data suggest that HFD-fed LKO livers possess altered cholesterol and bile acid metabolic pathways compared to WT mice, providing a mechanistic path to examine the resistance to DIO in LKO mice.

**Figure 3.**
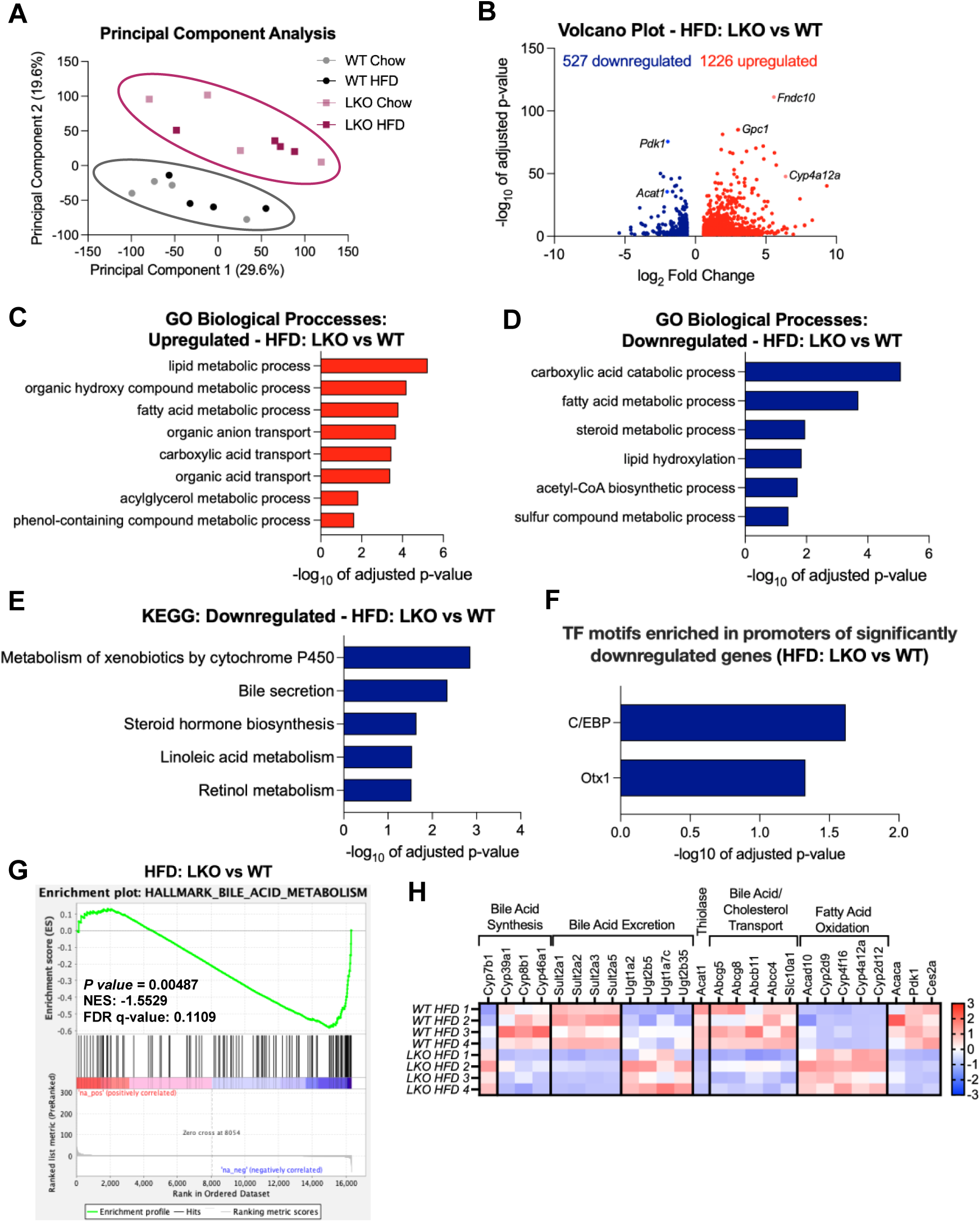
Transcriptional changes observed in female ARGLU1 LKO mice demonstrate decreased steroid and bile acid metabolism. Genes were significant if they passed an adjusted *P*-value of < 0.05 and log_2_ fold change ≥|1.5| as cut-offs. (A) Principal component analysis (PCA) of RNA-sequencing data demonstrates significant separation between WT and LKO genotypes. (B) Volcano plot of significantly different genes between LKO versus WT HFD-fed liver samples. Pathway enrichment analysis (g:profiler) was conducted for upregulated (C) and downregulated (D) biological pathways in HFD-fed LKO liver samples compared to WT controls. (E) KEGG analysis showed downregulated pathways in HFD-fed LKO mice compared to WT, with no significant upregulated pathways. (F) Transcription factor motifs enriched in promoters of either significantly downregulated or upregulated genes, comparing HFD-fed LKO mice to WT mice. (G) Gene-set enrichment plot comparing HFD-fed LKO mice to WT mice for bile acid metabolism. (H) Heatmap (z-score) of differentially expressed genes of interest, based on gene ontology. GO, gene ontology.

### HFD-fed LKO mice have decreased liver and plasma cholesterol compared to WT mice

Analysis of RNA-seq data highlighted distinct cholesterol and bile acid pathways that were different in the LKO mice compared to WT mice. Therefore, we assessed the levels of liver and plasma cholesterol in WT and LKO mice. Compared to WT, LKO mice had significantly reduced hepatic cholesterol (males: 51%; females: 84%; **Figure 4A**, **4H**) and plasma cholesterol levels (males: 40%; females: 26%; **Figure 4B**, **4I**). We next examined individual gene expression of key cholesterol synthesis and metabolic regulators in HFD-fed WT and LKO mice. Interestingly, there was no significant difference in cholesterol synthesis rate-limiting genes, *Hmgcs1* and *Hmgcr,* between female HFD-fed WT and LKO mice (**Figure 4C**); however, *Hmgcs1* was significantly decreased in male HFD-fed LKO mice compared to WT controls (**Figure 4J**), but this was not observed at the protein level (data not shown).

**Figure 4.**
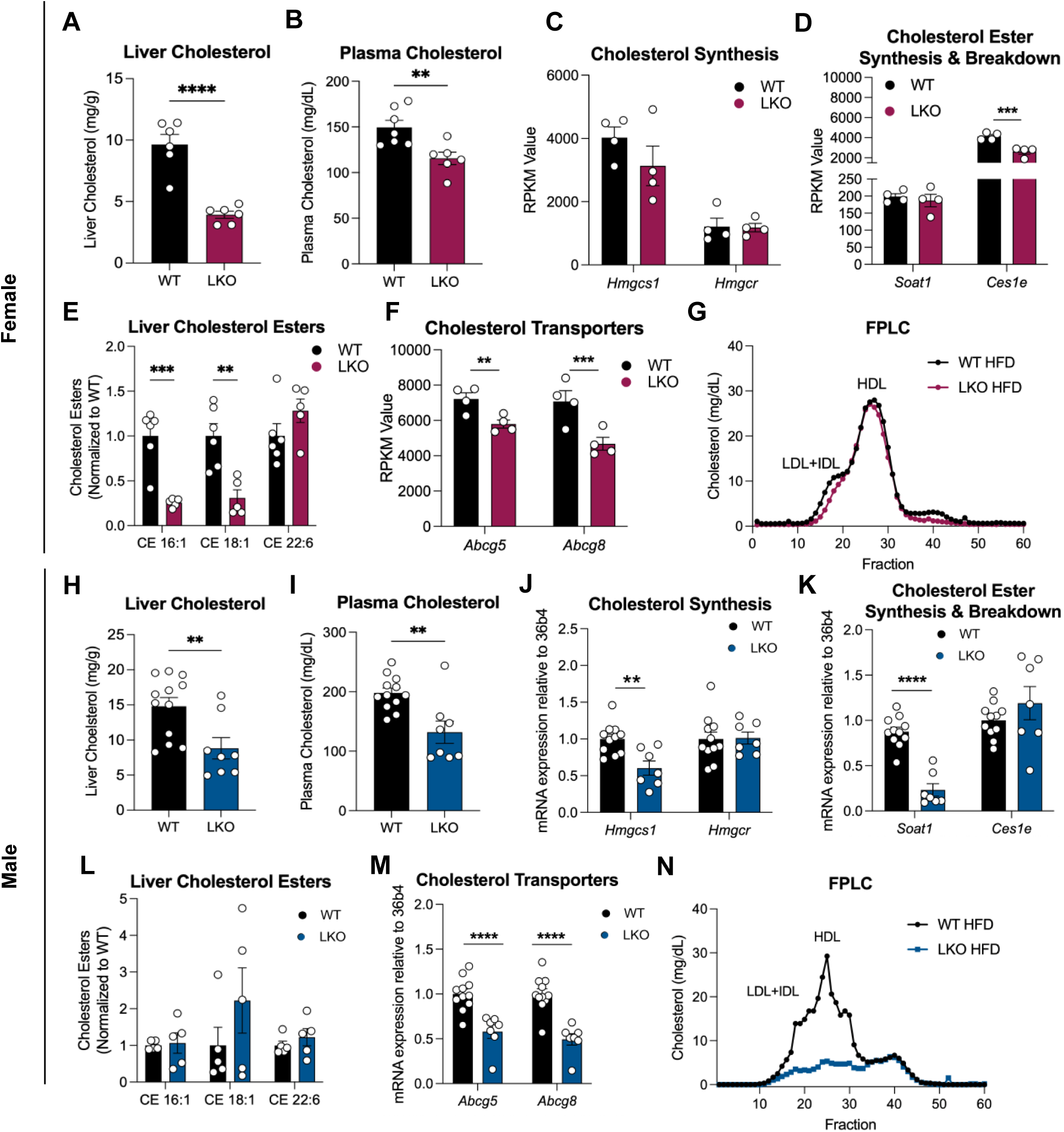
HFD-fed LKO mice have decreased hepatic and plasma cholesterol, not explained by changes in cholesterol synthesis, storage or export. Lipids were extracted from 12-week-HFD-fed female (A) or male (H) liver samples using the Folch method and assayed for cholesterol (N=6-12). Plasma collected from female (B) or male (I) mice (N=6-12) to measure plasma cholesterol through colorimetric assays. Transcriptomic analysis of genes involved in cholesterol synthesis (C, J), cholesterol ester synthesis and breakdown (D, K), and cholesterol transporters (F, M) (N=4-11). Quantification of cholesterol esterases from liver extracts by lipidomic analysis (LC/MS/MS) in female (E) and male (L) mice (N=5-6). FPLC from female (G) and male (N) HFD-fed WT and LKO plasma, pooled from four mice per genotype. \**P* ≤ 0.05, \*\**P* ≤ 0.01, \*\*\**P* ≤ 0.001, \*\*\*\**P* ≤ 0.0001, by unpaired-two tailed T-Test (A-F, H-M). Data are represented as mean ± SEM with individual animals noted as dots.

While *Soat1* (gene responsible for cholesterol ester synthesis) was not different between WT and LKO female mice (**Figure 4D**), *Ces1e,* a gene responsible for cholesterol ester catabolism,^21^ was decreased in female LKO mice compared to WT controls (**Figure 4D**).

Interestingly, we observed reduced liver cholesterol ester levels in female HFD-fed LKO mice compared to WT mice (**Figure 4E**). In male HFD-fed mice, LKO mice had decreased *Soat1* mRNA expression (**Figure 4K**) compared to WT mice, however, there was no difference between genotypes in *Ces1e* expression (**Figure 4K**) or liver cholesterol ester content (**Figure 4L**).

The mRNA expression of hepatic cholesterol efflux transporters (*Abcg5, Abcg8*) was markedly decreased in LKO mice compared to WT mice (**Figure 4F**, **4M**). No difference was observed in duodenum or jejunum cholesterol transporter expression between WT and LKO mice (**Figure S5A-B**). We performed FPLC to fractionate lipoproteins from pooled mouse plasma. Cholesterol-containing lipoproteins were reduced in all fractions in male HFD-fed LKO mice (**Figure 4N**) and were slightly lower in female HFD-fed LKO mice (**Figure 4G**) compared to WT control mice. Overall, these data suggest ARGLU1 has a role in cholesterol homeostatic pathways, but this role is not correlated with changes in cholesterol synthesis, export, or storage genes.

### LKO mice fed a HFD have decreased synthesis of 12α-hydroxylated bile acids compared to WT mice

Since bile acids are synthesized from cholesterol in the liver and are critical for lipid absorption, we investigated bile acid pathways in HFD-fed WT and LKO mice. LKO mice had decreased expression of hepatic bile acid synthesis (**Figure 5A, 5F**) and transporter (**Figure 5B, 5H**) genes. HFD-fed LKO mice showed a 65% (female) and 83% (male) decrease in *Cyp8b1* expression, a gene that encodes CYP8B1 which catalyzes the 12α-hydroxylation of bile acids (**Figure 5A, 5F**). By label free quantitative proteomics, CYP8B1 was detectable in WT samples, but below the limit of detection in LKO mice (**Figure 5G**). To assess if decreased CYP8B1 resulted in altered bile acid species, we measured individual bile acids from the bile collected from HFD-fed WT and LKO mice (**Table S4**). Interestingly, LKO mice had significantly decreased levels of the major 12α-hydroxylated bile acid, TCA, in both sexes compared to WT mice (**Figure 5C, 5I**).

**Figure 5.**
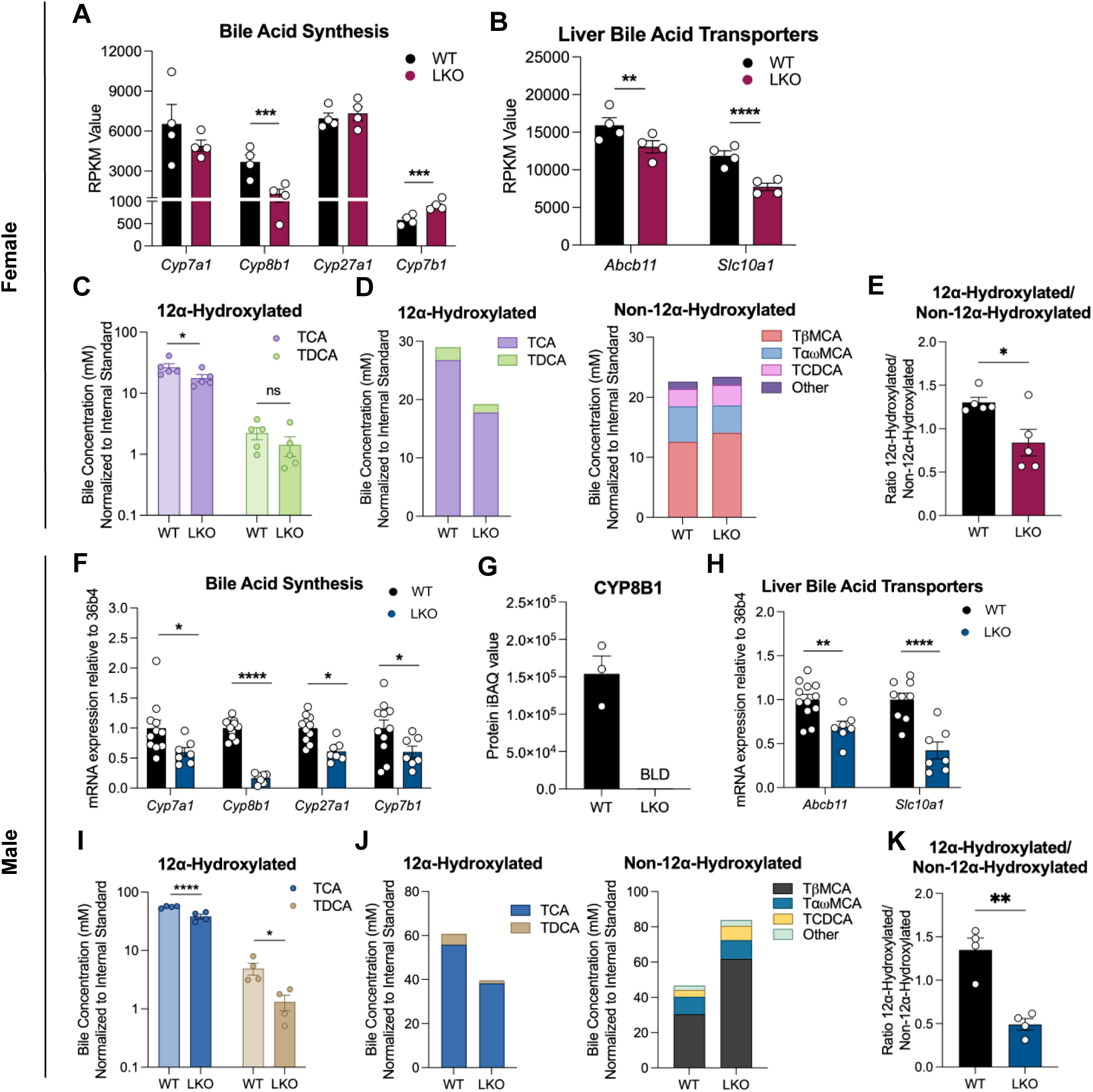
ARGLU1 LKO mice have decreased hepatic Cyp8b1 levels and 12α-hydroxylated bile acids. Hepatic transcriptomic analysis demonstrates decreased bile acid synthesis gene expression (A, F), and decreased bile transporter expression (B, H), (N=4-12). Bile was analyzed for 12α-hydroxylated bile acid species (TCA, TDCA) from female (C) and male (I) WT and LKO mice (N=4) (note log_10_ Y-axis). Total 12α-hydroxylated and non-12α-hydroxylated bile acids (D, J), and the ratio of the two groups was measured (E, K) (N=4). Proteomic analysis for CYP8B1 (G) from livers collected from male HFD-fed WT and LKO mice. BLD, below limit of detection; TCA, taurocholic acid; TDCA, taurodeoxycholic acid; TβMCA, Tauro-β muricholic acid; TαωMCA, Tauro-αω muricholic acid. \**P* ≤ 0.05, \*\**P* ≤ 0.01, \*\*\**P* ≤ 0.001, \*\*\*\**P* ≤ 0.0001, by unpaired-two tailed T-Test (A-K). Data are represented as mean ± SEM with individual animals noted as dots.

By measuring 12α-hydroxylated and non-12α-hydroxylated bile acid species in both sexes, (**Figure 5D, 5J**), we established that LKO mice had a significantly lower ratio of 12α-hydroxylated to non-12α-hydroxylated bile acids compared to WT mice (**Figure 5E, 5K**). A decrease in 12α-hydroxylated/non-12α-hydroxylated bile acids generates bile with increased hydrophilicity that is not as efficient at solubilizing fat and cholesterol.^13^ Analysis of fecal and liver samples showed decreased fecal CA (**Figure S6A, S6C, Figure S7**) and hepatic TCA (**Figure S6B, S6D**) content in LKO mice, compared to WT controls, consistent with the overall decrease in 12α-hydroxylated bile acids observed in bile of LKO mice. No difference was observed in ileal bile acid transporters (*Slc10a2*) between WT and LKO mice (**Figure S8A, S8B**). Collectively, these data suggest LKO mice have reduced hepatic *Cyp8b1* expression, corresponding to lower 12α-hydroxylated bile acid content in the bile, resulting in impaired dietary lipid absorption.

### Fatty acid oxidation is increased, and synthesis is decreased, in female LKO mice compared to control WT mice

The RNA-sequencing dataset revealed fatty acid metabolism as a significantly differentially regulated pathway in LKO mice compared to WT, thus we measured hepatic and plasma triglyceride content. Hepatic triglyceride levels were not different between WT and LKO mice of either sex (**Figure 6A**, **Figure S9A**). Plasma triglycerides were decreased by 38% (*P* = 0.04) in female LKO mice compared to WT (**Figure 6B**), while male LKO mice exhibited a 11% (*P* = 0.07) reduction compared to WT mice (**Figure S9A**). LKO mice on HFD had decreased expression of *de novo* lipogenic genes (**Figure 6C, Figure S9B**) and increased expression of fatty acid oxidation genes at the mRNA (**Figure 6D**, **Figure S9C**) and protein level (**Figure S9D**). Additionally, the lipid carboxylesterases *Ces2e* and *Ces2f,* markers of triglyceride hydrolysis, were increased in LKO mice compared to WT mice at the mRNA (**Figure 6E**) and protein level (**Figure S9E**), suggesting increased availability of fatty acids for oxidation. To assess hepatic VLDL secretion, we inhibited LPL activity with Poloxomer 407 and found no difference in hepatic VLDL secretion between female HFD-fed WT and LKO mice, indicating similar triglyceride production (**Figure 6F**). Lipidomic analysis of liver extracts revealed decreased diacylglycerols (DAG) for each carbon chain length (**Figure 6G**) and number of unsaturations (**Figure 6H**) in HFD-fed LKO compared to WT mice. Interestingly, FPLC demonstrated lower VLDL levels in female and male LKO mice compared to WT mice (**Figure 6I**, **Figure S9F**). The respiratory exchange ratio (RER), an indicator of fuel source used for energy, highlighted greater utilization of lipids as energy source in female HFD-fed LKO mice compared to WT mice (**Figure 6J**), a phenotype that was not statistically significant in males (**Figure S9G**). Collectively, these findings suggest that ARGLU1 has a role in lipid metabolism, with HFD-fed LKO mice exhibiting increased fatty acid oxidation, decreased triglyceride synthesis, and increased genes involved in triglyceride hydrolysis.

**Figure 6.**
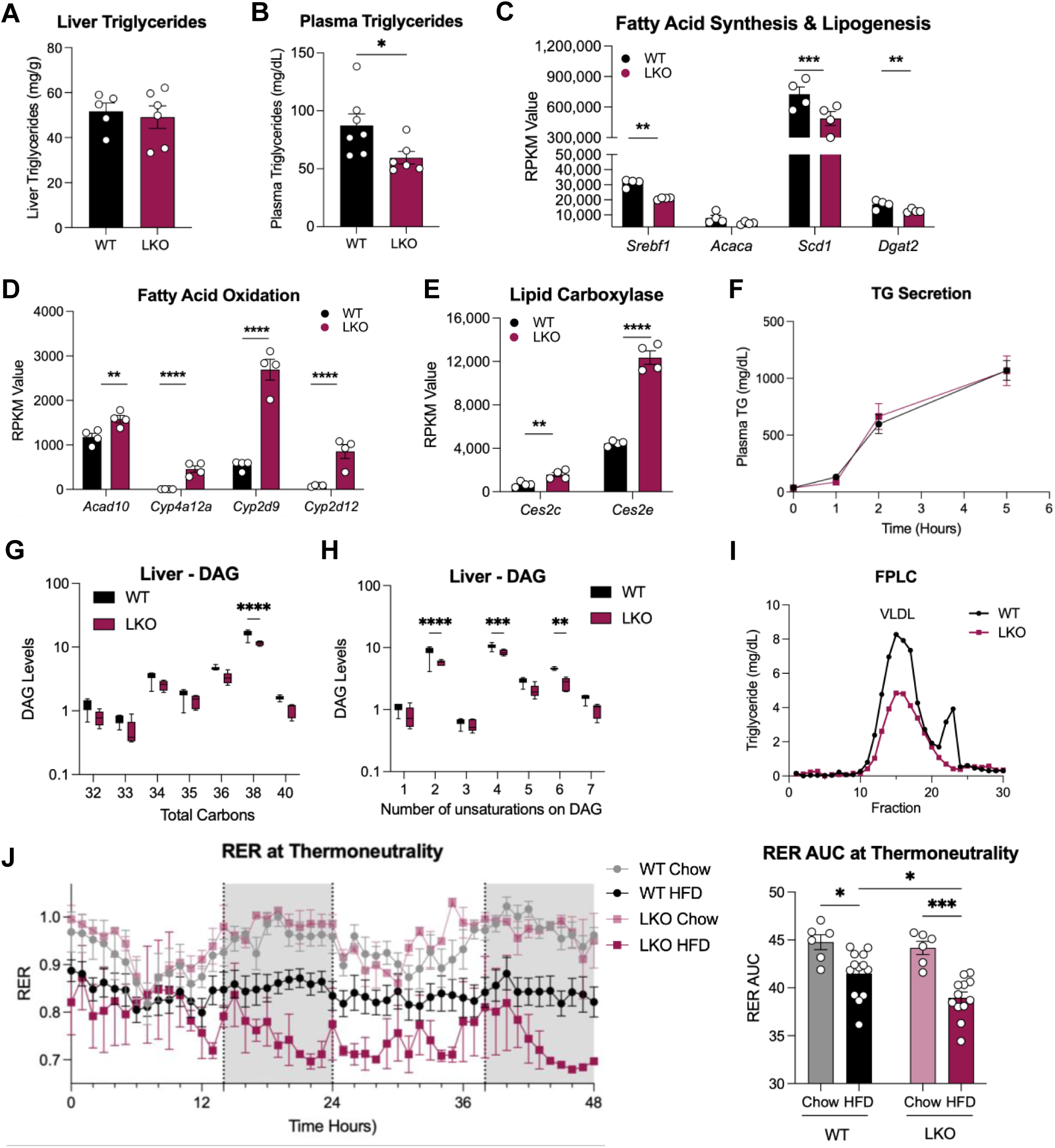
Increased expression of genes involved in fatty acid oxidation in female LKO mice compared to WT controls. Female WT and LKO mice fed HFD for 12-weeks. Liver triglyceride levels (A) and plasma triglycerides (B) from fed mice (N=4-7). RPKM values collected from HFD-fed WT and LKO mice fed or HFD diet for fatty acid synthesis and lipogenesis (C), fatty acid oxidation (D), lipid esterase genes (E) (N=4). (F) Triglyceride secretion from female mice injected i.p. with Poloxamer 407 at 1000 mg/kg in 0.9% NaCl (N=4-7). Lipidomic analysis of hepatic diacylglycerol (DAG) species from WT and LKO female mice (N=5) for total carbon (G) and saturation content (H). (I) FPLC fractionation of female HFD-fed WT and LKO mouse plasma pooled from four mice per genotype (fed state) and measured for triglyceride. (J) Female HFD- and chow-fed mice (N=6-7) were housed in Promethion Metabolic cages at thermoneutrality (30°C), from which oxygen consumption and carbon dioxide production were measured to analyze respiratory exchange ratio (RER). \**P* ≤ 0.05, \*\**P* ≤ 0.01, \*\*\**P* ≤ 0.001, \*\*\*\**P* ≤ 0.0001, by unpaired two-tailed T-Test (A-E) or two-way ANOVA followed by Holm-Sidak test (F-J). Data are represented as mean ± SEM with individual animals noted as dots.

### Protein interactome analysis of hepatic ARGLU1 in WT HFD-fed mice identifies TAF15 as binding partner

To identify *in vivo* binding partners of ARGLU1, we performed a Rapid Immunoprecipitation of Endogenous Proteins using Mass Spectrometry (RIME) study^22^ from female HFD-fed WT and LKO liver samples (**Figure S10A**). We identified 67 proteins enriched (≥1.0 log_2_ fold change) in WT versus ARGLU1 LKO livers (**Table S5**), including spliceosomal, transcription factor, and RNA processing proteins (**Figure S11**). From the ARGLU1 RIME data, the top five enriched proteins were ZC3H18, TAF15, PRPF38A, BUD31, NCBP2 (**Figure S10B**). These proteins all have roles in the spliceosome, with the exception of TAF15, which we investigated further. The RIME protein hits were also confirmed in our HEK293 ARGLU1-BioID experiment, performed previously,^18^ which identified ≥88% of the proteins recognized in our liver RIME dataset (≥1.0 log_2_ fold change), including TAF15 (**Figure S12**). TAF15 is a TATA-box binding protein associated factor and has been found to interact with C/EBP; and notably, the TAF15 liver knockout mouse has been reported to be protected against DIO.^23^ Interestingly, ChIP-sequencing datasets of C/EBP, HNF4α, and TAF15 in HepG2 cells demonstrate that C/EBP and HNF4α bind at the *CYP8B1* promoter region, with TAF15 and HNF4α binding 5kb upstream of the promoter in an area of open chromatin defined by H3K27ac (**Figure S13**). To explore if ARGLU1 can act as a coactivator for HNF4α, we performed a GAL4-hHNF4α/UAS luciferase reporter assay in HEK293 cells for HNF4α. The presence of ARGLU1 increased HNF4α activity by 1.74-fold (**Figure S14**). These data suggest that ARGLU1 could participate in transcription factor protein complexes (including ones containing TAF15, C/EBP, HNF4α) to provide selective regulation of metabolic gene expression (**Figure S10C**).

### Knockdown of ARGLU1 in mice with pre-existing obesity reduces weight gain

To address whether knockdown of ARGLU1 in liver could be a treatment for obesity, we performed an intervention study, in which WT mice with pre-existing DIO were injected with a virus to drive liver-specific Cre expression (**Figure 7A**). Injection of AAV8-Ttr-CRE into HFD-fed WT mice induced ARGLU1 knockdown (**Figure 7B**) and led to reduced body weight compared to GFP-injected controls (**Figure 7C-D**). Interestingly, the ARGLU1 knockdown group also had decreased hepatic expression of *Cyp8b1* (**Figure 7E**). Collectively, these findings demonstrate that knockdown of ARGLU1 can mitigate DIO.

**Figure 7.**
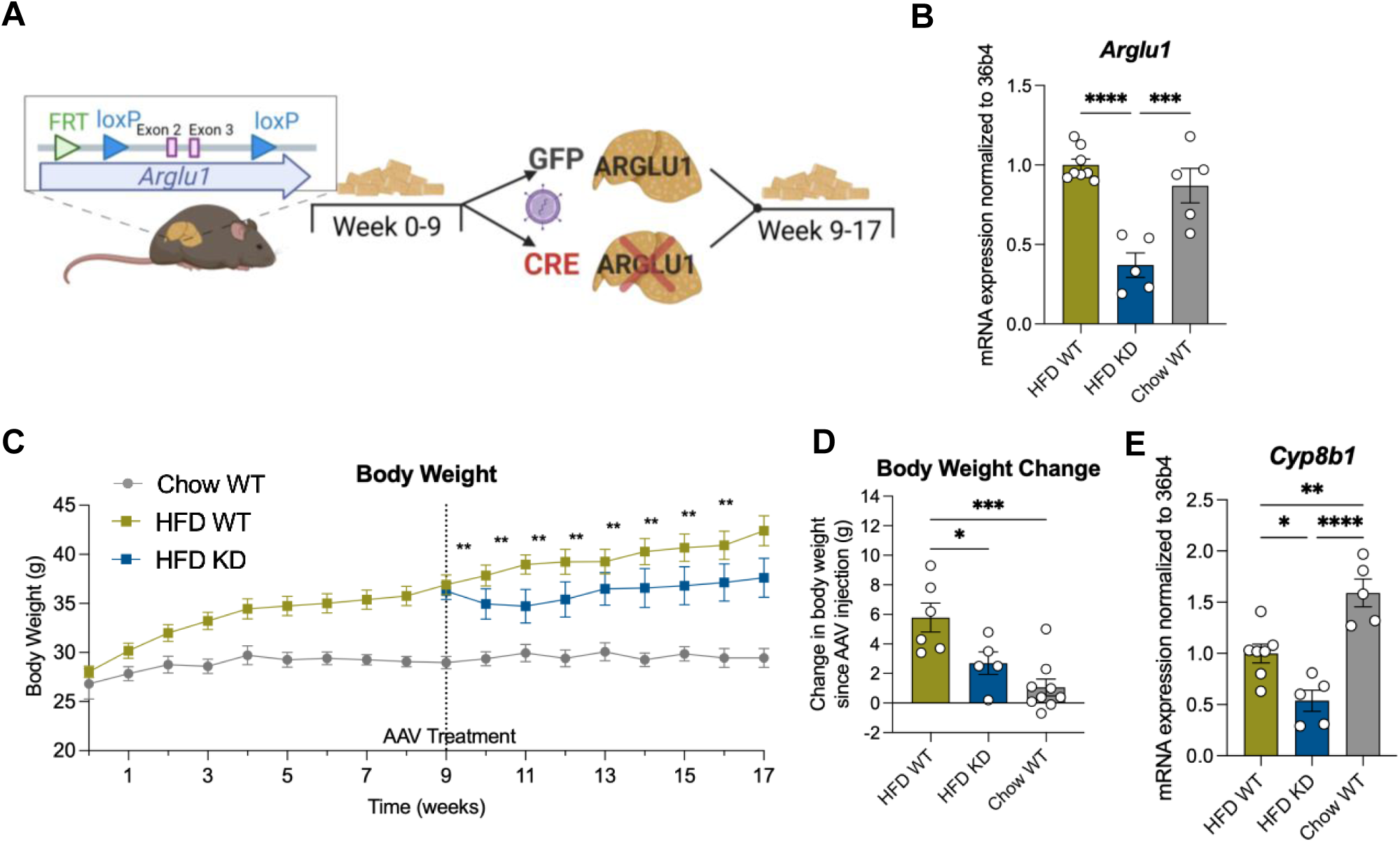
Knockdown of ARGLU1 in obese WT mice reduces weight gain. (A) Scheme of acute ARGLU1 viral knockdown experiment. (B) Hepatic gene expression of *Arglu1* at the end of study (week 17). (C) Body weight of male WT HFD-fed or chow-fed mice for 9 weeks, then IV-injected with either AAV8-Ttr-GFP or AAV8-Ttr-CRE and continued on respective diet for 8 additional weeks. (D) Body weight gain difference from AAV-injection until the end of the study. (E) Hepatic *Cyp8b1* mRNA expression. \**P* ≤ 0.05, \*\**P* ≤ 0.01, \*\*\**P* ≤ 0.001, \*\*\*\**P* ≤ 0.0001, by two-way ANOVA followed by Holm-Sidak test (B-E). Data are represented as mean ± SEM with individual animals noted as dots (N=5-7).

## DISCUSSION

The growing prevalence of metabolic diseases worldwide highlights the urgent need for novel anti-obesity therapies. The new incretin-based anti-obesity therapies act centrally to decrease food intake and, while extremely effective, they can have adverse side effects that contribute to discontinuation. However, there are myriad metabolic pathways that provide opportunities for pharmacologic intervention. Herein, we provide evidence that the transcriptional coregulator protein ARGLU1 contributes to DIO by regulating bile acid synthesis and consequently lipid absorption.

The absence of ARGLU1 in hepatocytes was sufficient for male and female mice to be resistant to DIO without changes to food intake, energy expenditure, or locomotion. We observed increased excretion of fecal cholesterol and triglycerides in LKO mice. In line with this, we found that LKO mice had reduced biliary 12α-hydroxylated bile acids (synthesized by CYP8B1) compared to WT mice, thereby altering the hydrophilicity of bile and its capacity to emulsify dietary lipids. The LKO mice appeared healthy and were indistinguishable from WT mice, apart from body weight. Notably, the colour and physical appearance of the feces collected from WT and LKO mice were comparable, suggesting that LKO mice do not have overt steatorrhea. We surmise that LKO mice are did not exhibit malabsorption of necessary dietary lipids and fat-soluble vitamins, since LKO mice still gained weight on HFD compared to chow-fed LKO mice. In addition, there were minimal differences in the basal body weights between chow-fed WT and LKO mice.

Our findings of resistance to DIO and decreased hydrophobic 12α-hydroxylated bile acids, are similar to those reported in *Cyp8b1*^-/-^ mice.^13,17^ After 4 weeks of HFD (TD. 88137, same diet used in our study), male *Cyp8b1^-/-^* mice weighed 15% less than WT mice. *Cyp8b1^-/-^* mice were reported to have decreased lipid absorption, with >3.5-fold increase in fecal triglycerides, and 43% lower plasma [^3^H]-triolein levels compared to WT mice.^13^ This decrease in plasma [^3^H]-triolein levels was reversed with the re-introduction of TCA in *Cyp8b1^-/-^* mice,^13^ highlighting the importance of bile acid composition in lipid absorption. *Cyp8b1^-/-^*female mice also showed improved lipid tolerance (30% decrease in AUC of lipid tolerance test),^24^ reduced adipose tissue weights,^24^ and improved glucose homeostasis^24,25^ compared to WT mice. Cholesterol absorption is also impaired with altered bile acid composition. For example, male *Cyp8b1^-/-^* mice had 40-50% decreased cholesterol absorption.^17^ Male *Cyp8b1^-/-^* mice had an 80% decrease in hepatic cholesterol,^13^ findings comparable to ARGLU1 LKO mice (60% decrease). Notably, human heterozygous carriers of CYP8B1 complete loss-of-function mutations were more insulin sensitive, and had reduced 12α-hydroxylated to non-12α-hydroxylated bile acid ratio in plasma.^26^ Given the myriad beneficial effects of decreased CYP8B1 activity, several pharmaceutical companies (Merck, Xenon) and academic laboratories have patented CYP8B1 inhibitors for the treatment of dyslipidemia, atherosclerosis, metabolic dysfunction-associated steatotic liver disease (MASLD) and diabetes.^27-32^

LKO mice also exhibited distinct lipid metabolism, including reduced hepatic expression of *Dgat2*, a lipogenic regulator,^33^ and increased expression of *Ces2e*, a triglyceride hydrolysis gene,^34^ suggesting LKO mice have decreased triglyceride synthesis and increased triglyceride hydrolysis. Mice housed in metabolic cages displayed lower RER values, indicating increased lipid utilization in female LKO mice. These findings indicate that any lipids absorbed in LKO mice are preferentially oxidized rather than stored, contributing a second mechanism by which LKO mice are resistant to DIO.

We had previously identified ARGLU1 as a coregulator of a variety of nuclear receptors;^18^ and expected to see some interplay with HFD-feeding. However, following the analyses of RNA-seq data, no specific nuclear receptor involvement was apparent. To investigate the molecular mechanism underlying ARGLU1’s effects, we conducted a RIME study to identify ARGLU1-interacting proteins, and found RNA-binding and transcriptional regulatory proteins, such as TAF15. Hepatic TAF15 KO mice demonstrate resistance to DIO to a similar magnitude as ARGLU1 KO mice when placed on the same diet, with ∼25% decrease in weight gain and ∼50% decrease in visceral adipose tissue weight.^23^ Furthermore, TAF15 has been reported to regulate nuclear translocation of C/EBP,^23^ a transcription factor that regulates adipogenesis, gluconeogenesis and amino acid catabolism.^35^ More recent findings suggest that C/EBP plays a role in cholesterol metabolism^36^ and can regulate farnesoid X receptor (FXR) *in vitro*, a transcription factor involved in bile acid metabolism.^37^ Additionally, C/EBPβ^-/-^ mice were resistant to DIO body weight gain when fed a 60% HFD, and also exhibited reduced hepatic lipogenic gene expression.^38,39^ ARGLU1 BioID data generated previously using HEK293 cells^18^ found ARGLU1 interacted with both TAF15 and C/EBP in these cells, suggesting a potential molecular mechanism. Examination of ChIP-sequencing data for TAF15 and C/EBP from HepG2 cells, we found that C/EBP binds to DNA at the *CYP8B1* promoter, and TAF15 binds 5Kb upstream in a region of open chromatin (defined by H3K27ac).

To assess if targeting ARGLU1 could be a model for treatment intervention, we performed an ARGLU1 knockdown experiment in DIO WT mice, where the knockdown of ARGLU1 resulted in weight loss compared to WT mice. This finding provides support for the therapeutic targeting of ARGLU1, where selective ablation of hepatic ARGLU1 could decrease obesity. Several small molecules have been developed to target the interaction between a transcription factor and a transcriptional coregulator, providing a tractable mechanism to increase transcriptomic specificity.^40-42^

In conclusion, our study indicates that loss of hepatic ARGLU1 can attenuate HFD-induced obesity. We found that ARGLU1 promotes lipid absorption and therefore impacts overall lipid metabolism. Due to the conserved regulation of bile acid synthesis between mouse and human, inhibition of hepatic ARGLU1 may be a potential target for the treatment of obesity.

### Limitations of the study

The main limitation is that we have yet to define at a molecular level the interaction partners of ARGLU1 at specific sites on the chromatin. Regulation of CYP8B1 is critical for the phenotype and should be examined at a molecular level by ChIP and/or heterologous promoter reporter assays to assess whether ARGLU1 can co-activate TAF15, HNF4α and/or C/EBP on the CYP8B1 promoter/enhancer. Another limitation is that we observed increased fatty acid oxidation from indirect calorimetry and attribute this to the liver using primarily RNA-seq/qPCR data, but we cannot rule out contributions from other tissues.

Although most of our studies were performed in both sexes, RNA-seq and global proteomic analysis were only completed in one sex each. Future experiments could address this by assessing chow- and HFD-fed WT and LKO liver samples for RNA-seq in male mice and proteomic analysis in female mice.

## RESOURCE AVAILABILITY

### Lead contact

Requests for further information and resources should be directed to and will be fulfilled by the lead contact, Carolyn Cummins.

### Materials availability

ARGLU1 LKO mice generated in this study are available from the lead contact with a completed materials transfer agreement.

### Data and code availability

This paper does not report original code. All newly generated datasets have been deposited, and identifiers are available in the key resources table.

This paper analyzes existing, publicly available, accessible ChIP-Seq data. The reference for these data is listed in the key resources table.

Any additional information required to reanalyze the data reported in this paper is available from the lead contact upon request.

## ACKNOWLEDGMENTS

We gratefully acknowledge financial support from the Canadian Institutes of Health Research (PJT-156194, PJT-198086 to C.L.C.) and Canadian Foundation for Innovation (#33088 to C.L.C.); and stipend support from Pfizer Canada Graduate Fellowship in Pharmaceutical Sciences (S.B.C.), Canada Graduate Scholarship (S.B.C.), and Ontario Graduate Scholarship (S.B.C.). We thank Vlera Fonda and Henriette Uhlenhaut for sharing the tissue-specific RIME protocol, and Rafael Montenegro-Burke and K. Sandy Pang for their gifts of bile acid standards. We also would like to acknowledge all past and current Cummins Lab trainees that assisted with this project. In particular, we thank Florian Le Billan and Zhi Di (Judy) Deng for processing the liver samples used in the proteomic analysis. The graphical abstract and other figure schemes were generated with BioRender.com.

## AUTHOR CONTRIBUTIONS

Conceptualization, S.B.C., L.M., and C.L.C; formal analysis, S.B.C., M.F.S., J.S., C.K.; funding acquisition, C.L.C.; investigation S.B.C., L.M., S.A.S., F.M., M.F.S., J.S., P.E.T., C.K., T.K.; methodology S.B.C., L.M., M.F.S., A.A., C.L.C.; resources S.A.L., P.E.T.; supervision, S.A., A.A., J.L.B., C.L.C.; visualization, S.B.C.; writing—original draft, S.B.C., C.L.C.; writing—review & editing, all authors.

## SUPPLEMENTAL INFORMATION

Document S1. Figures S1–S14 and Tables S1, S2, S4, S6.

Table S3. RNA-sequencing GO downregulated and upregulated Biological Pathways, comparing HFD-fed LKO mice to HFD-fed WT mice, related to Figure 3.

Table S5. Proteins identified from RIME using liver from HFD-fed WT and LKO mice, related to Figure S10, S11, and S12.

## STAR★METHODS

### KEY RESOURCES TABLE

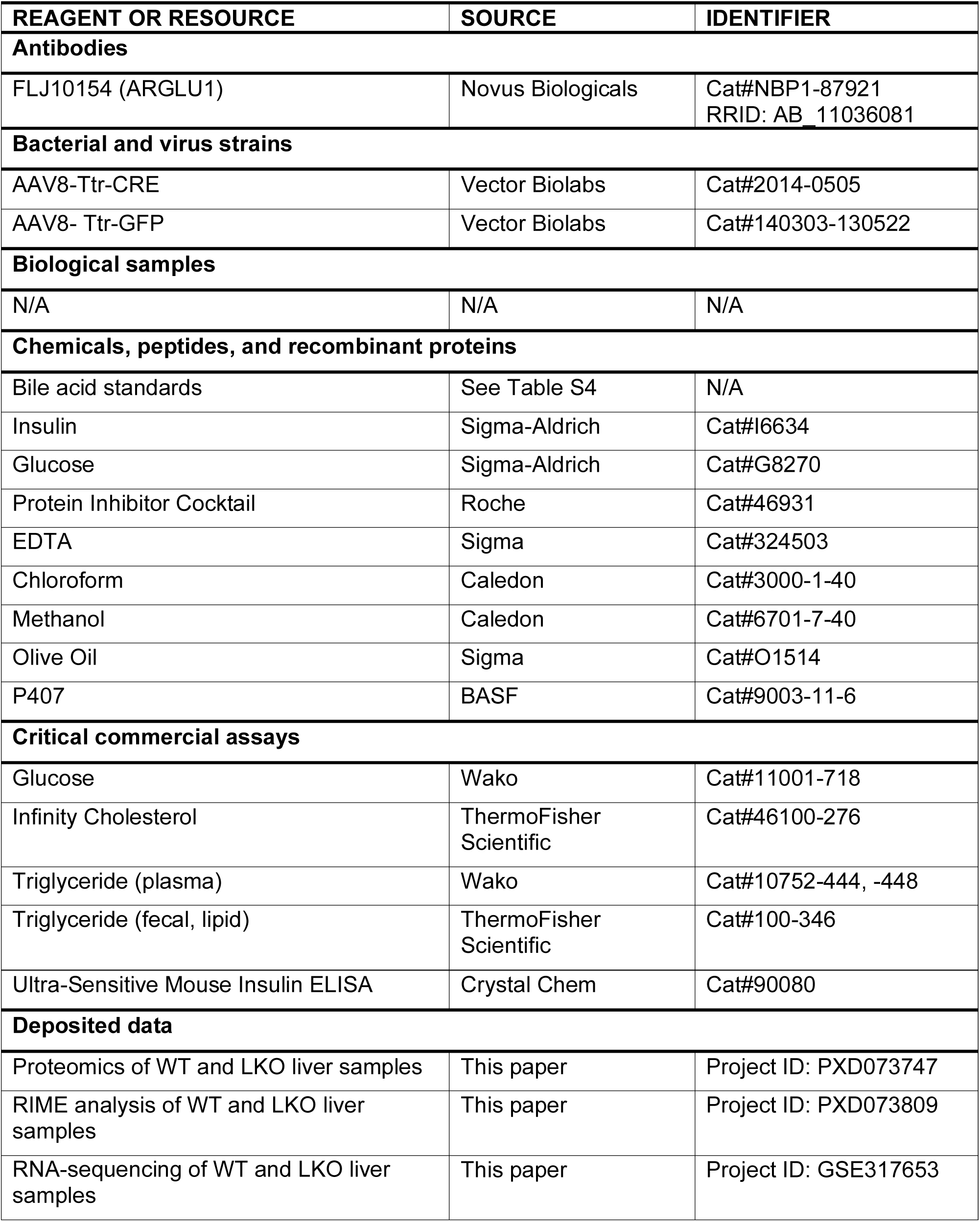
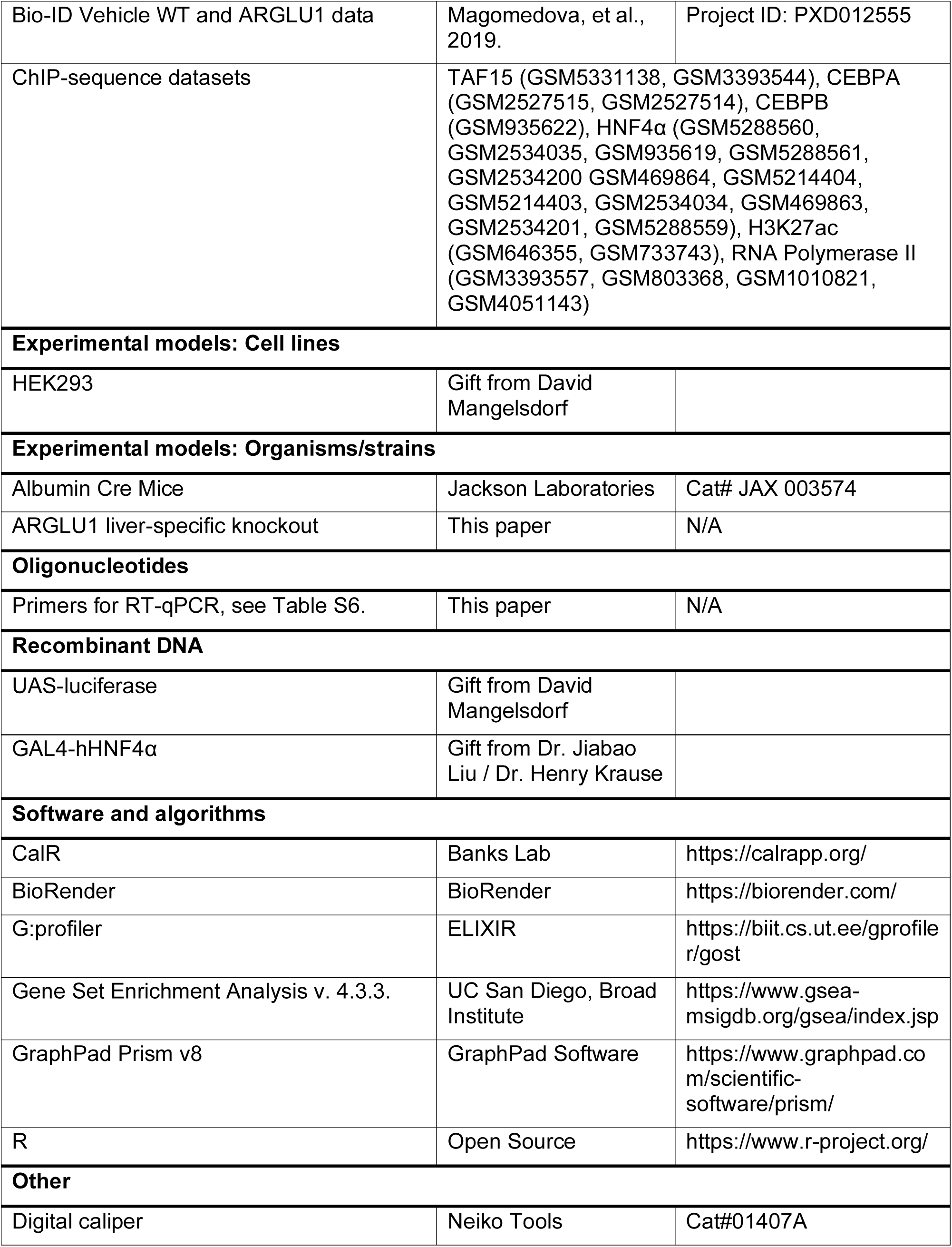
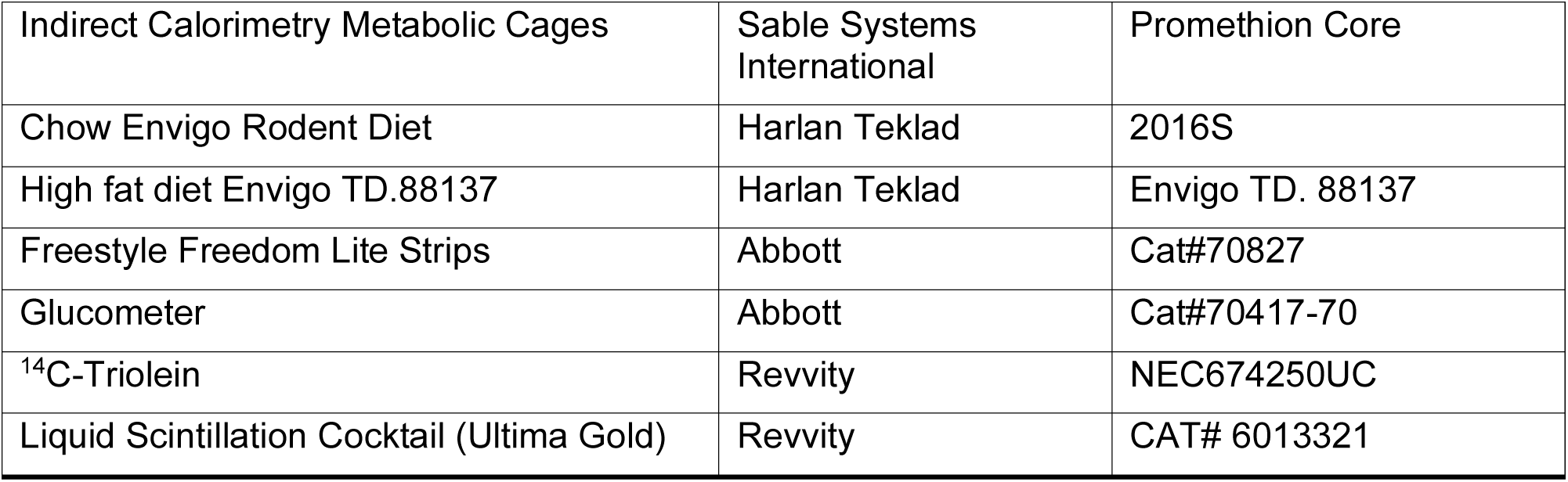

#### Luciferase reporter assay

Human embryonic kidney (HEK293) cells were cultured in Dulbecco’s modified Eagle’s medium (DMEM) supplemented with 10% fetal bovine serum (FBS). Cells were seeded 3.5 x 10^4^ cells per well of 96-well plate in DMEM containing 10% charcoal-stripped FBS. DNA was transfected using calcium-phosphate, as previously described,^18^ with total plasmid DNA content (150 ng) consisting of 50 ng of UAS-luciferase reporter, 20 ng of β-galactosidase, 15 ng of HNF4α LBD of interest (GAL4-hHNF4α), and either 15 ng of CMX vector, CMX-hARGLU1. Twenty-four hours post transfection, luciferase values were read and normalized to β-galactosidase to control for transfection efficiency and expressed as normalized luciferase output.^5^

#### Animal studies

Mice were bred and housed in the Division of Comparative Medicine at the University of Toronto, within a standard temperature and light-controlled environment. All mice in experiments were littermate controls *Arglu1^fl/fl^*wild-type (WT) and *Arglu1^fl/fl^* Albumin Cre liver-specific knockout mice (LKO) age-matched between ages 5-months and 10-months old. No difference in body weight gain was observed between WT and Albumin Cre controls (Albumin^Cre/+^) (**Table S2**), and so *Arglu1^fl/fl^* mice were used as WT controls in experiments. Both male and female mice were used in experiments and mice were bred and reared on a standard rodent chow (Envigo, 2016S rodent diet, Harlan Teklad Mississauga, ON). All protocols were approved by the Faculty of Medicine and Pharmacy Animal Care Committees at the University of Toronto (Toronto, ON).

For diet-induced obesity (DIO) experiments, mice were fed a high-fat, high-cholesterol diet (HFD) containing 42% kcal from fat and 0.2% cholesterol (Envigo, TD.88137, Harlan Teklad Mississauga, ON) for 12 weeks (**Table S1**). Experimental mice were either singly or doubly housed. Body weight was measured weekly, and food intake was measured biweekly. Animals were sacrificed by decapitation at 10 a.m. Trunk blood was collected in tubes containing 5 μL of 0.5 M EDTA, with plasma isolated by centrifugation at 500 x g, 4°C for 20 minutes. Tissue samples were weighed, aliquoted and snap frozen in liquid nitrogen and stored at -80°C. To assess body size from which to normalize tissue weight, the right femur from each mouse was isolated and cleaned, and length measured using a digital caliper (Neiko Tools, Taiwan).

For the induced knockdown of ARGLU1, WT mice were injected with 2x10^11^ viral gc/mouse AAV8-Ttr-Cre (Cat#2014-0505) or AAV8-Ttr-GFP (Cat#140303-130522, Vector Biolabs, Malvern, PA) for controls in 0.9% NaCl delivered via tail-vein injection after 9 weeks of HFD or chow feeding. Post-injection body weight was measured for an additional 8 weeks on the respective diet.

#### Measurement of fat content by DEXA scan

Body composition of WT and LKO mice on chow or HFD was measured using dual-energy X-ray absorptiometry (DEXA, Bruker). Mice were anesthetized with 2.5% isoflurane-oxygen, placed in a prone position and limbs and head were extended from the body. X-ray images were taken at two energy levels to determine hard tissues (lean and bone), and total tissues (lean, bone, and fat tissues) by using 0.08 mm AI and 0.0 mm AI filters, respectively. Images were corrected for illumination and converted to density units. In defining the region of interest (ROI), the head was excluded. To calculate the total fat percent the following equation was used:

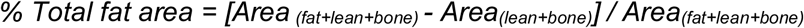

To measure visceral fat mass, the ROI was defined as a box spanning L1/L2 vertebrae through L4/L5 vertebral space.

#### Indirect calorimetry

Mice were individually housed in temperature controlled Promethion Core Metabolic cages (Sable Systems International, Las Vegas, NV) in the Department of Comparative Medicine at the University of Toronto. Mice had free access to food and water with cage enrichment. After an 8-week period on chow or HFD, mice were place in the Promethion metabolic cages either at room temperature or thermoneutrality of 30°C for a 3-day acclimation period prior to 5-day data collection. The final 48 hours were used to calculate basal and total whole body energy expenditure (volume O_2_ consumption and CO_2_ production), respiratory exchange ratio (RER), locomotor activity, food intake, and water consumption.

#### Fecal lipid absorption assay

Mice were individually housed with fresh bedding for a 72-hour period in the final week of the study. Stools were separated from bedding, dried over a 3-day period in a fume hood, and homogenized. Lipids were extracted using the Folch method, using 2:1 v/v chloroform/methanol solution. The extracted lipid content from the diets and stools was assessed gravimetrically.

Next, intestinal lipid absorption was calculated from total lipid output (based on fecal weight and lipid content) and dietary lipid consumed over 72-hours.

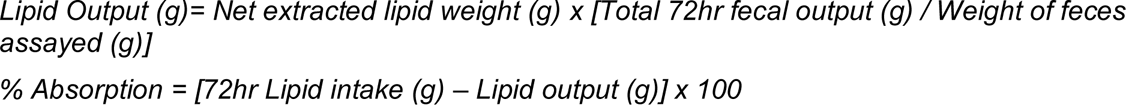

#### Glucose and insulin tolerance tests

During week 9 of HFD feeding, a glucose tolerance test (GTT) was performed after a 16-hour fast with *ad libitum* access to water. Local anesthetic was applied at the tail (EMLA cream, AstraZeneca, Cambridge, UK). After basal blood glucose collection (20 µL) into EDTA-coated microvette tubes, 20% D-glucose prepared in 0.9% NaCl (at 10 μL/g of lean body weight) was delivered i.p., and blood glucose levels were monitored at 15, 30, 60, 120 minutes, from the tail using a glucometer (Abbott, Chicago, IL) and test strips (Freestyle Freedom Lite, Alameda, CA). After 11 weeks on HFD, an insulin tolerance test (ITT) was performed where mice were fasted for 4 hours, and basal tail-vein blood glucose was collected using a glucometer (Abbott).

Female mice were dosed i.p. at 0.75 U/insulin/kg of lean body weight and males were dosed i.p. at 1 U/insulin/kg of lean body weight. Blood glucose was measured 15, 30, 45, 60, 90 minutes post insulin administration.

#### Plasma metabolite analyses

After decapitation, trunk blood was collected into EDTA-coated tubes and placed on ice. After centrifugation at 500 x g, 4°C for 20 minutes, plasma was isolated and stored at -80°C. Plasma triglycerides (Wako, Richmond, VA), cholesterol (ThermoFisher), and glucose (Wako) were measured by colorimetric assays. Plasma insulin was measured from fed and overnight fasted male mice using an Ultra-Sensitive Mouse Insulin ELISA kit (Crystal Chem, Elk Grove Village, IL).

#### Liver lipid analysis

Lipids were extracted from liver samples (100 mg) using the Folch method as previously described ^43,44^. Briefly, liver was homogenized in 4 mL of chloroform/methanol (2:1, v/v), washed with 50 mM NaCl, and centrifuged at 1500 x g for 30 minutes. The organic phase was transferred to new tubes and washed twice in 0.36 M CaCl_2_/methanol, followed by centrifugation at 1500 x g for 10 minutes. The organic phase was transferred into 5 mL volumetric flasks and topped up to 5 mL with chloroform. Aliquots of standards and samples were dried down and dissolved in 10 µL of 1:1 chloroform/Triton X-100, and quantified for triglycerides (Infinity, ThermoFisher Scientific) and cholesterol (ThermoFisher Scientific) using colourimetric reagents.

#### RNA isolation, cDNA synthesis, and real-time qPCR analysis

From liver and intestinal tissues, total RNA was extracted using TRIzol® Reagent (ThermoFisher Scientific). For each sample, 2 µg RNA was incubated with DNase I and reverse transcribed into cDNA by random hexamer primers and using the High-Capacity Reverse Transcription System (Applied Biosystems (ABI), Burlington, ON). Reat-time quantitative PCR (qPCR) reactions were performed in 384-well plates, using CFX384 Real-time System C1000 Touch Thermal Cycler instrument (Bio-Rad; Bio-Rad, Hercules, CA). Each well contained 12.5 ng cDNA, 150 nM of each forward and reverse primers (**Table S3**), and 5 μL 2xSYBR Green PCR Master Mix (ABI). Using the comparative Ct method, relative mRNA levels were calculated normalizing to *36b4* mRNA.

#### RNA-sequence analysis

Total RNA was extracted from liver samples collected from female WT and LKO mice fed chow or HFD for 12 weeks using TRIzol® (ThermoFisher). Samples were processed for quality control, mRNA library preparation with Poly A enrichment by NEBNext Ultra II RNA, and sequencing of 20 million NovaSeq Paired-End150 reads by Illumina (Novogene, Sacramento, CA). Fastq files were provided, and analysis was performed using a previously optimized workflow.^45^ Sequence reads were aligned to the mouse reference genome (mm10) using Bowtie2. DESeq2 analysis was used to determine differentially expressed genes between the conditions. Genes were significant if they passed an adjusted *P*-value of ≤ 0.05 and log_2_ fold change ≥ |1.5| as cut-offs. Pathway enrichment analysis was conducted using g:profiler, a tool that identifies biological processes based on gene ontology terms.^46^ Gene set enrichment analysis was conducted following previous methodology.^47,48^

#### VLDL secretion assay

Mice were anesthetized locally at the tail using EMLA cream and injected i.p. with poloxamer 407 (p407; BASF, Mississauga, ON) at 1000 mg/kg in 0.9% NaCl. Blood was collected into EDTA-coated microvette tubes at 0, 1, 2, 5 hours post-injection. Blood was centrifuged at 500 x g for 20 minutes at 4°C to isolate plasma for triglyceride analysis.

#### Lipoprotein separation by Fast Protein Lipid Chromatography (FPLC)

Trunk blood was collected into EDTA-coated tubes and plasma was isolated through centrifugation at 500 x g for 20 minutes at 4°C. An NGC FPLC (Bio-Rad) equipped with an Enrich SEC 650 10 x 300 mm column was used. The running buffer consisted of 20 mM monosodium phosphate, 31 mM disodium phosphate, 150 mM NaCl at pH 7.0. Plasma samples from each group were pooled (65 µL/mouse was pooled to reach a total volume of 240 µL) and loaded into the system at a constant flow of 1 mL/min. Fractionation was started after 23 minutes and 250 µl of diluted plasma was collected for each fraction. These fractions were then analyzed using triglyceride and cholesterol assays, as described in previous sections.

#### Bile acid quantification

Bile acids were extracted from liver tissue and fecal samples according to previously reported methods.^49^ Flash-frozen liver samples (100 mg) were homogenized in 4 mL methanol spiked with CDCA-d4 10 µM (10 µL). After an overnight incubation at room temperature, samples were centrifuged at 8000 x g for 10 minutes, 90 µL of the supernatant was collected into glass vials spiked with TCA-d4 10 µM (10µL) for LC-MS/MS analysis. Fecal samples (10 pellets) were lyophilized overnight and homogenized. A total of 5 mg of fecal sample powder was resuspended in 1 mL MeOH and was spiked with CDCA-d4 10 µM (10 µL). Fecal samples were sonicated and centrifuged at 12000 x g for 10 minutes, where 90 µL of the supernatant was collected into glass vials and was spiked with TCA-d4 10 µM (10 µL). Samples were stored at - 20°C until run on the LC-MS/MS. Bile samples were prepared from 1 µL of bile collected from the gall bladder and immediately diluted 1:10 in methanol. Each diluted bile sample (195 µL) was combined with 5 µL of 8 µM TCA-d4 as an internal standard.

Standard solutions of individual bile acids (**Table S4**) were prepared by dissolving the compounds separately in methanol and used to generate a 5-point calibration curve.^49^ For each calibration point, internal standard TCA-d4 10 µM was used to quantify bile acid content of samples, while CDCA-d4 was used to confirm efficient sample extraction.

#### Liver proteomic analysis

Hepatic protein was extracted from male WT and LKO mice fed chow or HFD. Analysis was conducted as previously described.^18,50^ Livers were lysed with RIPA buffer and 100 μg of protein was digested using the FASP method on 30 kDa spin filters (Millipore).^51^ Eluted peptides were acidified and desalted using in-house made C18 pipette tips (10 μg capacity). Analysis was performed on an Easy nLC-1200 coupled to a QExactive HF mass spectrometer (ThermoFisher Scientific) operating in top 20 mode. The mobile phase was composed of Buffer A (0.1% formic acid) and Buffer B (0.1% formic acid in 80% acetonitrile). Peptides were separated using a PepMap RSLC C18 2 μm, 75 μm × 50 cm column and a PepMap 100 C18 3 μm, 75 μm × 2 cm precolumn with a 2-hour gradient of 5%–40% Buffer B. Data were analyzed using MaxQuant (v1.6.10.43)^52^, Perseus^53^, and MetaboMiNR^54^. The mass spectrometry proteomics data have been deposited to the ProteomeXchange Consortium via the PRIDE^55^ partner repository with the dataset identifier PXD073747.

#### Liver lipidomic analysis

Lipids were extracted from liver samples (100 mg) using the Folch method as previously described^43,44^ and dried under the fume hood. Samples were reconstituted in a 4:3:1 mixture of isopropanol, acetonitrile, and water. To 100 µL of sample, 5 µL of SPLASH LIPIDOMIX (Avanti Research) was added as an internal standard. A QC sample was prepared by mixing an equal volume of each sample together to form an average sample.

The samples were then processed on a Thermo Vanquish UPLC coupled to a Thermo Exploris 240 mass spectrometer. The separation was done on a Water Acquity BEH C18 column (100Å, 1.7 µm, 2.1-mm x 100 mm) at 55°C. The mobile phase was composed of a 40:60 (v/v) mix of water and acetonitrile with 0.1% formic acid and 10 mM ammonium formate (Buffer A) and a 90:10 (v/v) mix of isopropanol and acetonitrile with 0.1% formic acid and 10 mM ammonium formate (Buffer B). The gradient was as follows: 0 min (0% B), 5 min (50% B), 17.5 min (100% B), 18.5 min (100% B), 20 min (0% B), 22 min (0% B) at 400 µL/min. The data were collected in both positive and negative mode using a high resolution (120,000) MS1 scan for quantitation.

For lipid identification, the QC samples were processed in both polarities using a smart data dependent acquisition mode by the AcquireX software from Thermo Scientific. Peak identification and lipid identification were done using MS-DIAL5.^54,56^

#### Lipid tolerance test

Female and male HFD-fed WT and LKO mice were fasted overnight, and basal blood sample was collected. Mice were administered 200 µL olive oil (Sigma) by oral gavage, and 50 µL blood was collected from the tail using EDTA-coated microvette tubes at 1, 3, and 5 hour-post oral gavage.^20^ Plasma was isolated, and triglycerides were measured using a colorimetric triglyceride assay (Wako).

#### Triglyceride absorption assay

After 4-hour fast (7am-11am), each mouse was administered 1 µCi of ^14^C-Triolein (NEC674250UC, Revvity, Waltham, MA) dissolved in 200 µL olive oil via oral gavage, as previously described.^57,58^ Mice were sacrificed two hours post-gavage, and 100 µL of plasma was combined with 3 mL of Liquid Scintillation Cocktail (Revvity) prior to counting on a liquid scintillation counter.

#### Rapid immunoprecipitation of endogenous proteins using mass spectrometry (RIME)

This study followed methodology previously described^59^ on liver samples from female WT and LKO mice fed a HFD for 12 weeks. Samples underwent an overnight immunoprecipitation with 6 µg of ARGLU1 antibody (FLJ10154, Novus, Chesterfield, MO), or a no antibody control. Raw mass spectrometry data were first processed in MaxQuant using the integrated Andromeda search engine against the UniProt mouse proteome database. The resulting proteinGroups.txt output was imported into Perseus for downstream analysis. Label-free quantification intensity values were log_2_-transformed, and proteins with valid values were retained for analysis. Missing values were imputed from a normal distribution to account for signals from low-abundance proteins. The list of proteins was then normalized to determine the difference between WT and LKO detected proteins. The protein list was run through STRING Database Web-based app to determine protein-protein interaction clustering (https://string-db.org).^60^ Overlapping and unique proteins were identified using Venn diagrams (http://bioinfomatics.psb.ugent.be) comparing the RIME dataset against a previously reported *in vitro* HEK293 BioID ARGLU1 protein interactome.^18^ The mass spectrometry proteomics data have been deposited to the ProteomeXchange Consortium via the PRIDE^55^ partner repository with the dataset identifier PXD073809.

#### Quantification and statistical analysis

Data are presented as mean ± SEM. Outliers were identified with Grubb’s test with alpha=0.2. The number of animals/samples is indicated in the figure legends. Significance was calculated using two-way ANOVA followed by Holm-Sidak for multiple comparisons or an unpaired two-tailed T-Test when comparing only two groups. An ANCOVA was performed to determine difference between energy expenditure regression plots. Data were analysed using GraphPad Prism 10.2 Software (GraphPad, San Diego, CA). *P* ≤ 0.05 was considered statistically significant.

