## Supplemental File for "Liver-specific loss of the transcriptional coregulator ARGLU1 protects against diet-induced obesity in mice through decreased lipid absorption"

**DOCUMENT S1**

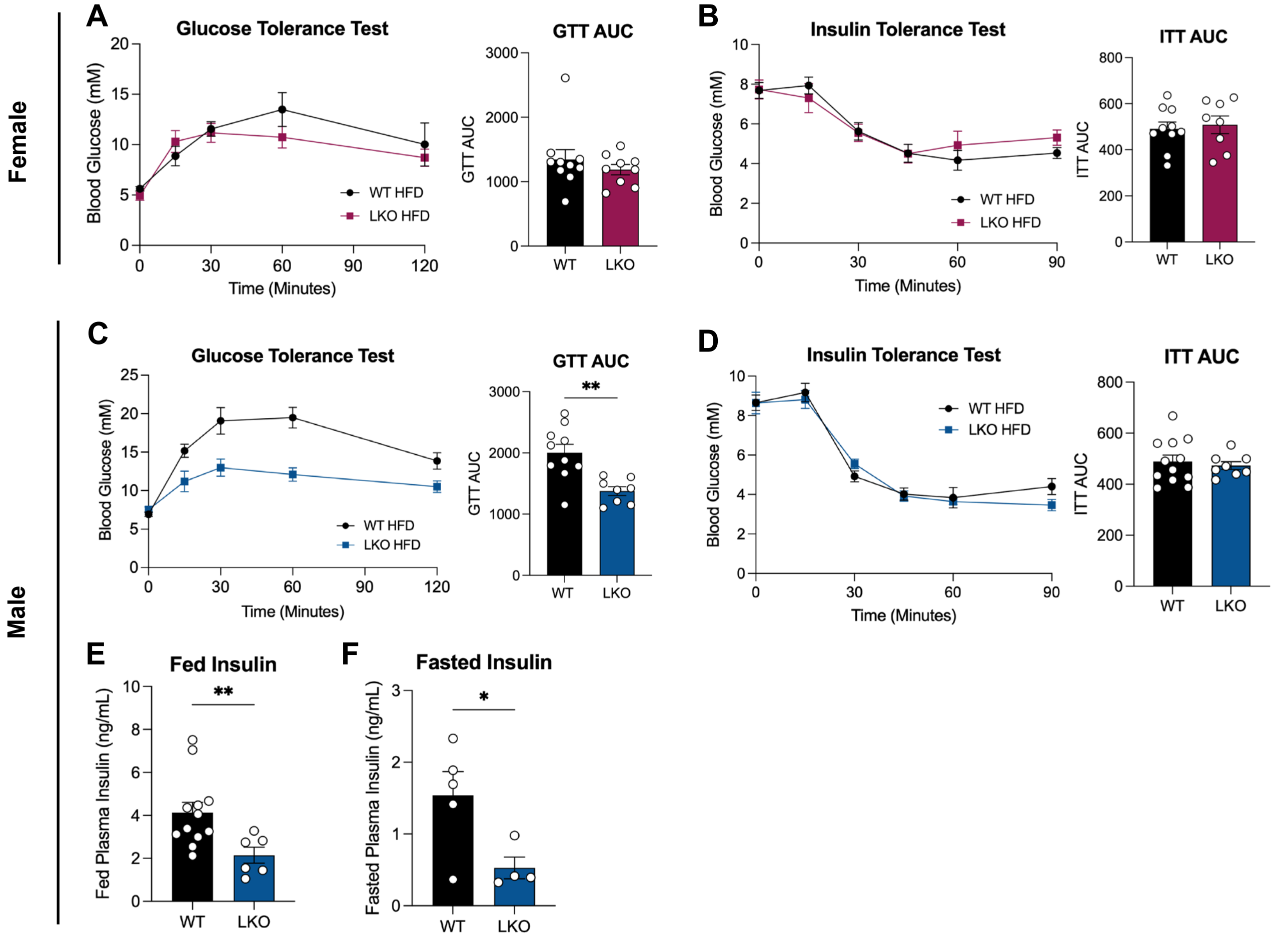


**Figure S1. LKO male mice are more glucose tolerant than WT mice on a HFD.** At 9 weeks of HFD-feeding, a glucose tolerance test (GTT) was performed in overnight fasted female (A) or male (C) mice, with 20% D-glucose prepared in 0.9% NaCl (at 10 µL/g of lean body weight). After a 4-hour fast, an insulin tolerance test (ITT) was performed in female (dosed at 0.75 U/insulin/kg of lean body weight i.p.) (B) or male (dosed at 1U/insulin/kg of lean body weight i.p.) (D) mice. Plasma insulin from fed (E) and fasted (F) male mice. **P* ≤ 0.05, ***P* ≤ 0.01, ****P* ≤ 0.001, *****P* ≤ 0.0001, by unpaired T-Test (E, F, AUC analysis). Data are represented as mean ± SEM with individual animals noted as dots (N=4-12).

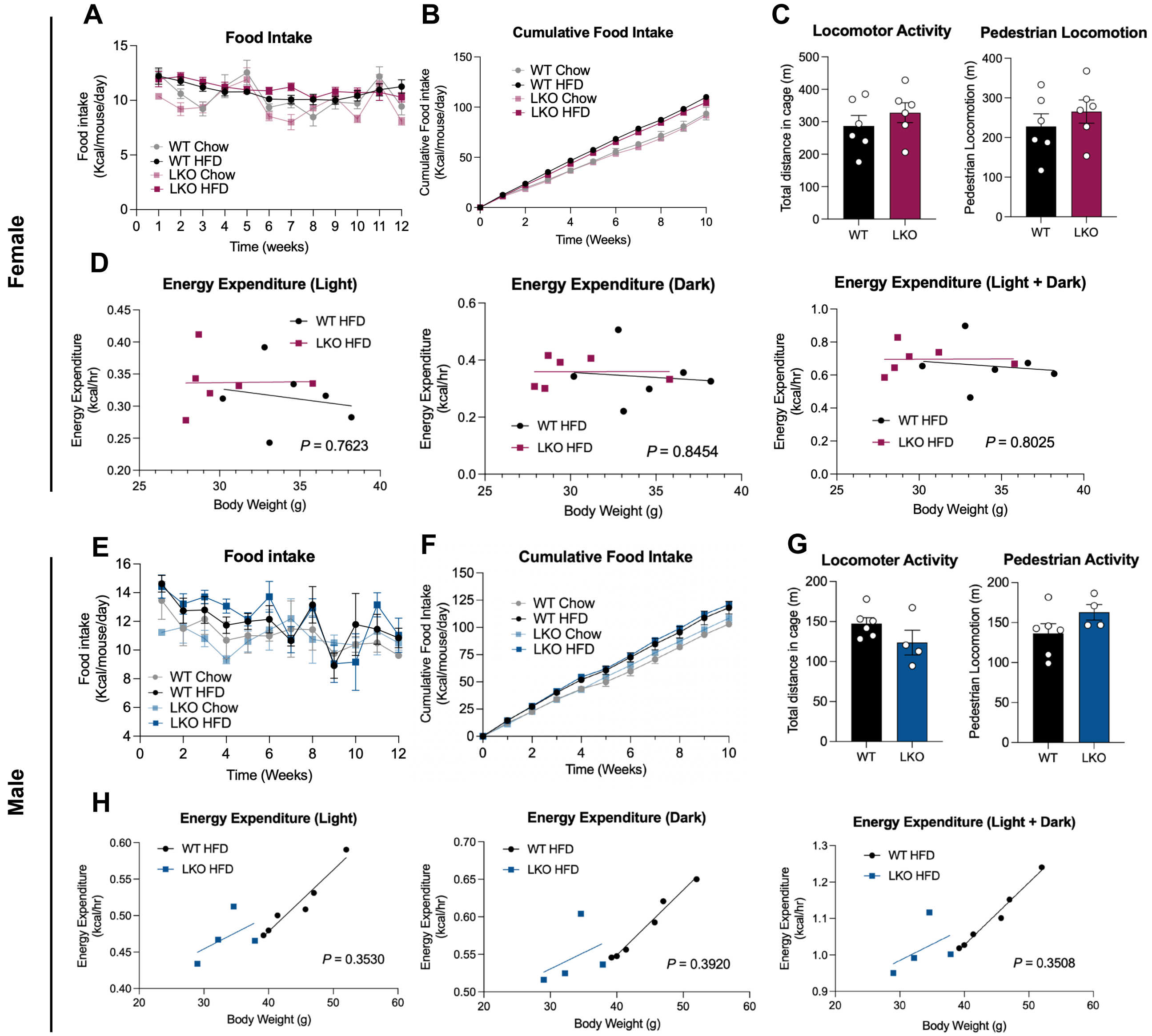


**Figure S2. Resistance to DIO in LKO mice is not related to food intake, locomotion, or energy expenditure.** Daily (A, E) and cumulative (B, F) food intake was measured biweekly for both female and male mice. Using Promethion metabolic cages, locomotor activity and pedestrian activity was measured for female (C) and male (G) mice after 11 weeks on HFD. Analysis of energy expenditure during light, dark and combined phases was measured for female (D) and male (H) mice analyzed using ANCOVA with body weight as a variable. *P* values represent differences between genotypes. **P* ≤ 0.05, ***P* ≤ 0.01, ****P* ≤ 0.001, *****P* ≤ 0.0001, by unpaired two-tailed T-Test. Data are represented as mean ± SEM with individual animals noted as dots (N=4-17).

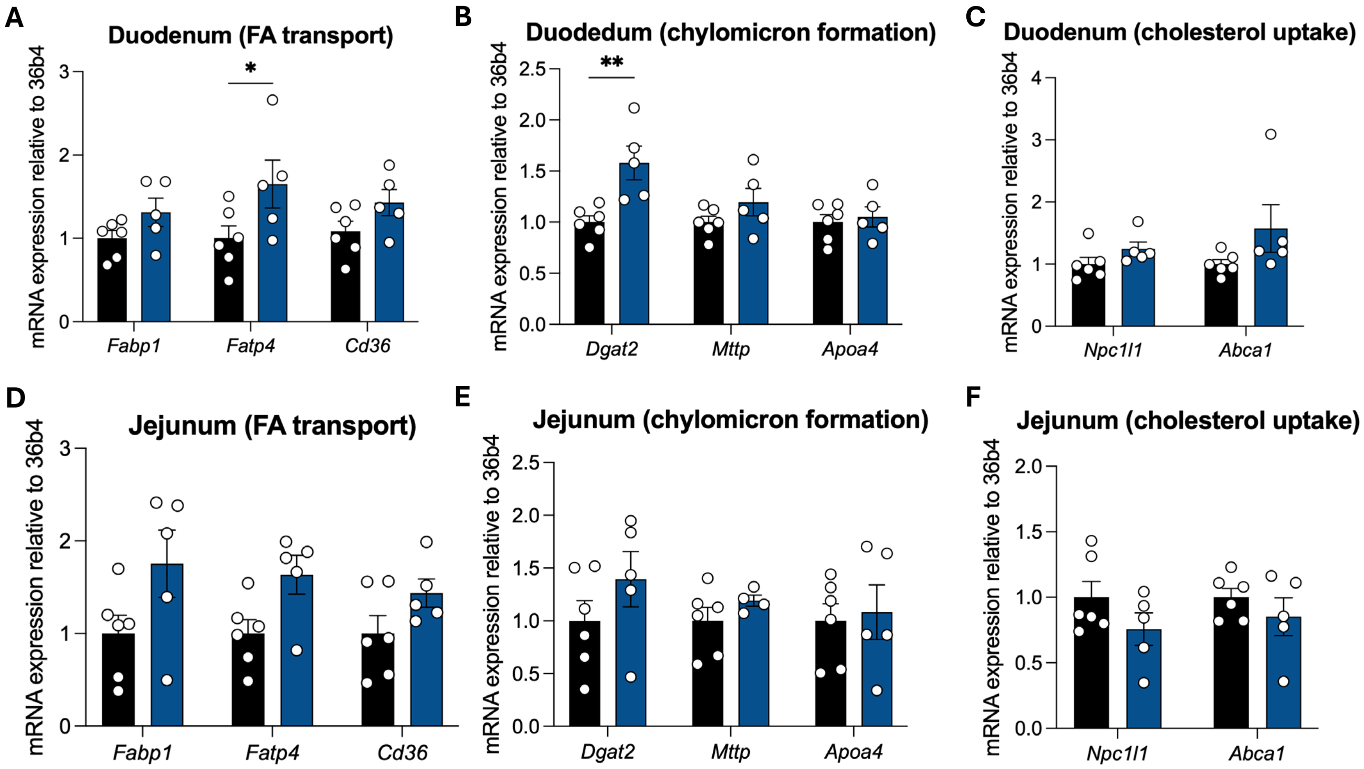


**Figure S3. No difference in intestinal fatty acid transporter, chylomicron formation, or cholesterol uptake transporter gene expression between WT and LKO mice.** Duodenum (A, B, C) and jejunum (D, E, F) samples were isolated from HFD-fed male mice for assessment of fatty acid (FA) transport and chylomicron formation gene expression. **P* ≤ 0.05, ***P* ≤ 0.01, ****P* ≤ 0.001, *****P* ≤ 0.0001, by unpaired two-tailed T-Test. Data are represented as mean ± SEM with individual animals noted as dots (N=5-7).

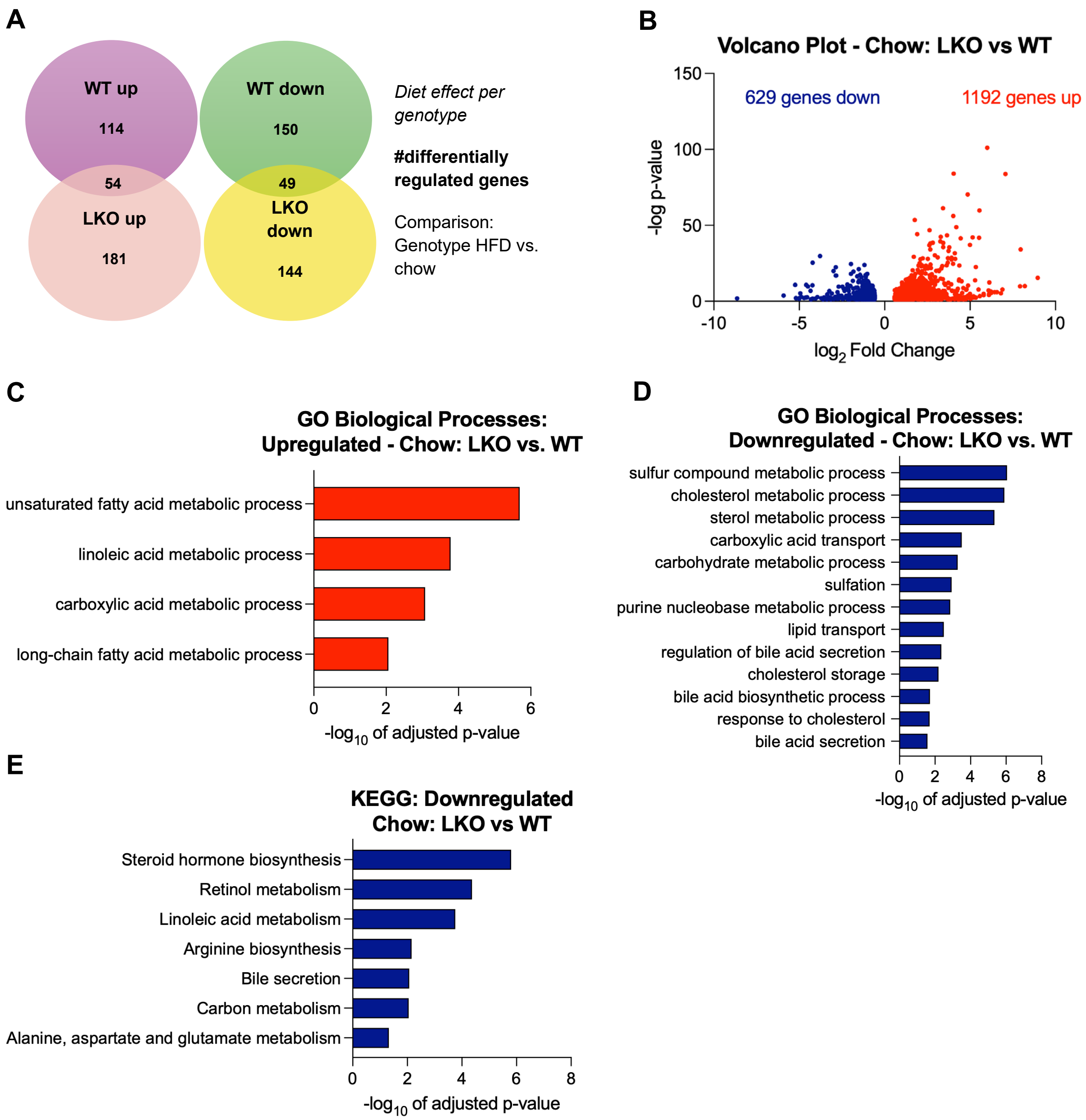


**Figure S4. Significant transcriptional differences between WT and LKO chow-fed female mice.** Pathway enrichment analysis was conducted using g:profiler, a tool that identifies biological processes based on gene ontology terms. (A) Venn diagram of differentially expressed genes for HFD versus chow for each genotype, comparing shared upregulated and downregulated genes for female WT and LKO mice. (B) Volcano plot of significantly different genes between LKO versus WT chow-fed liver samples. Upregulated (C) and downregulated (D) gene ontology (GO) pathways in chow-fed LKO liver samples compared to WT controls. (E) Downregulated KEGG pathway analysis on chow-fed LKO vs WT liver samples.

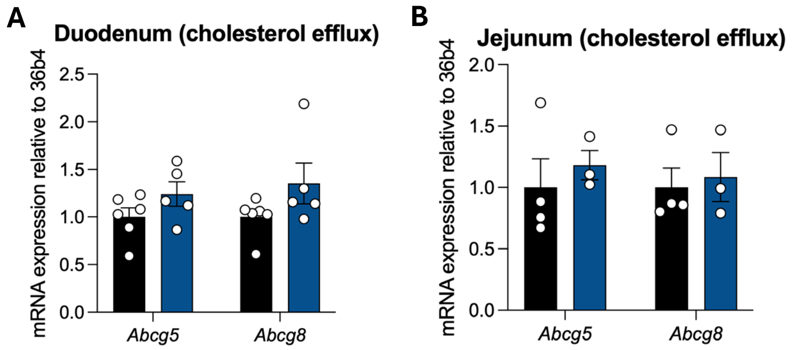


**Figure S5. No difference in intestinal cholesterol efflux transporter gene expression between WT and LKO mice.** Duodenum (A) and jejunum (B) samples were isolated from male mice for cholesterol transporter gene expression analysis. *$P \leq 0.05$, **$P \leq 0.01$, ***$P \leq 0.001$, ****$P \leq 0.0001$, by unpaired two-tailed T-Test. Data are represented as mean ± SEM with individual animals noted as dots (N=3-6).

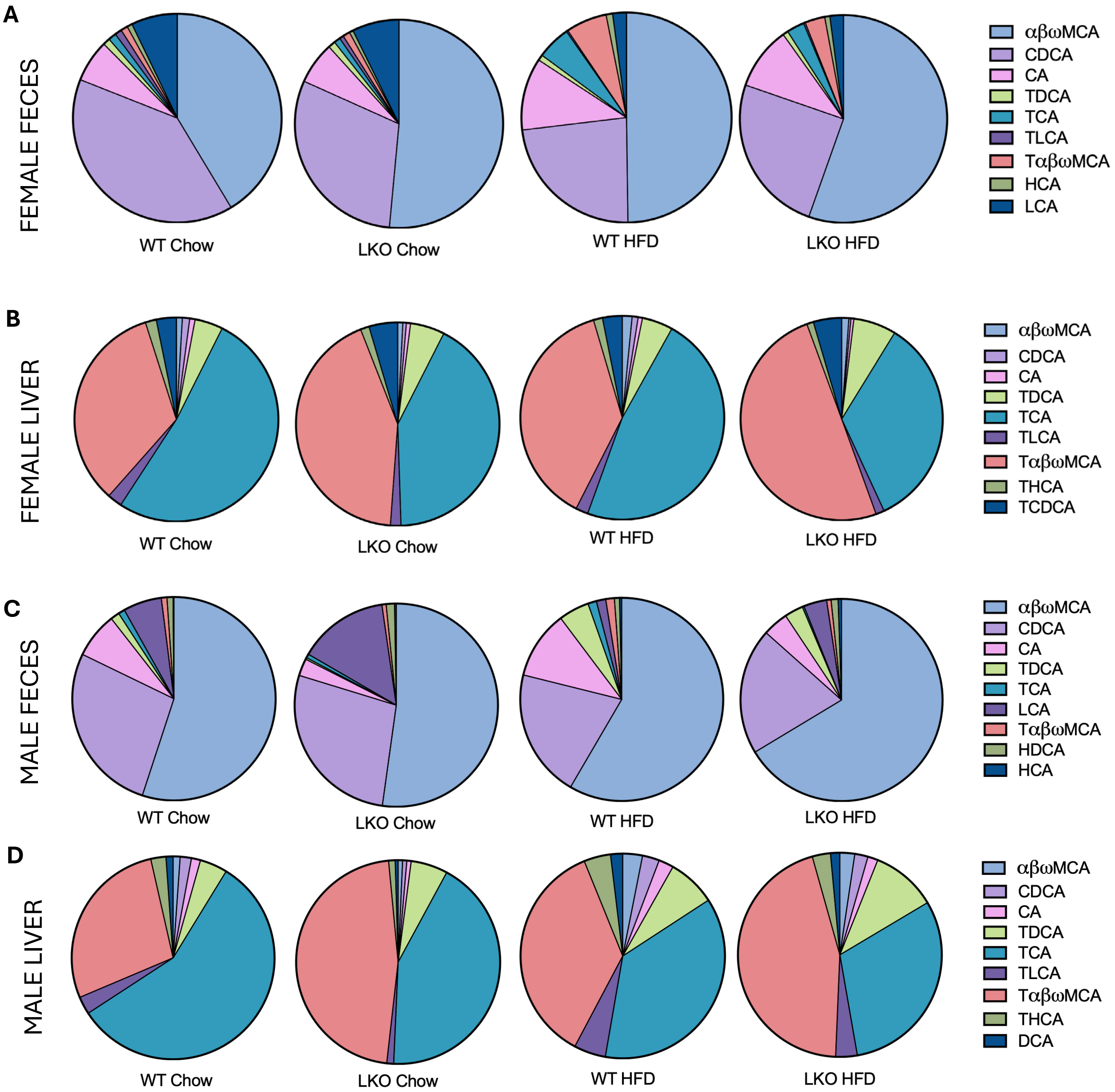


**Figure S6. Altered bile acid composition in hepatic and fecal samples from LKO mice compared to WT mice.** Ratio of bile acid content from female feces (A), female liver (B), male feces (C), male liver (D) samples for WT and LKO chow- and HFD-fed mice.

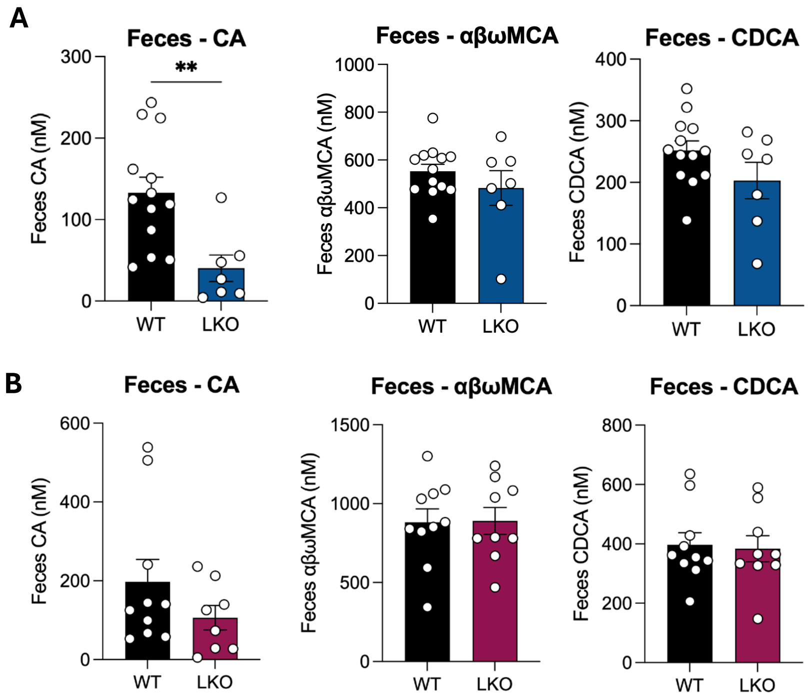


**Figure S7. Male LKO mice have decreased fecal cholic acid content compared to WT mice.** Fecal bile acid content from male (A) and female (B) HFD-fed mice, reported as nM. $*P \leq 0.05$, $**P \leq 0.01$, $***P \leq 0.001$, $****P \leq 0.0001$, by unpaired two-tailed T-Test. Data are represented as mean ± SEM with individual animals noted as dots (N=7-13).

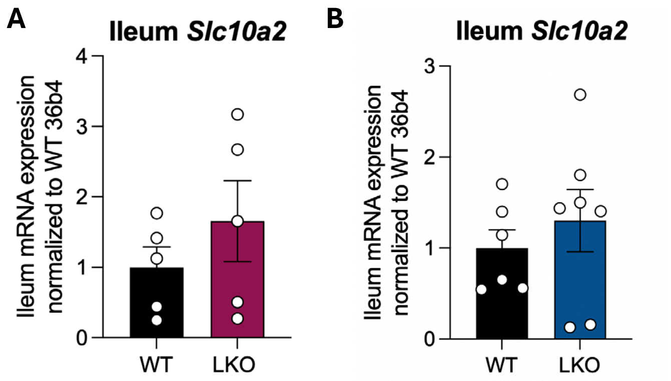


**Figure S8. WT and LKO mice have similar ileal bile acid transporter expression.** Expression of *Slc10a2* in ileum from female (A) and male (B) HFD-fed WT and LKO mice. *$P \leq 0.05$, **$P \leq 0.01$, ***$P \leq 0.001$, ****$P \leq 0.0001$, by unpaired two-tailed T-Test. Data are represented as mean ± SEM with individual animals noted as dots (N=5-7).

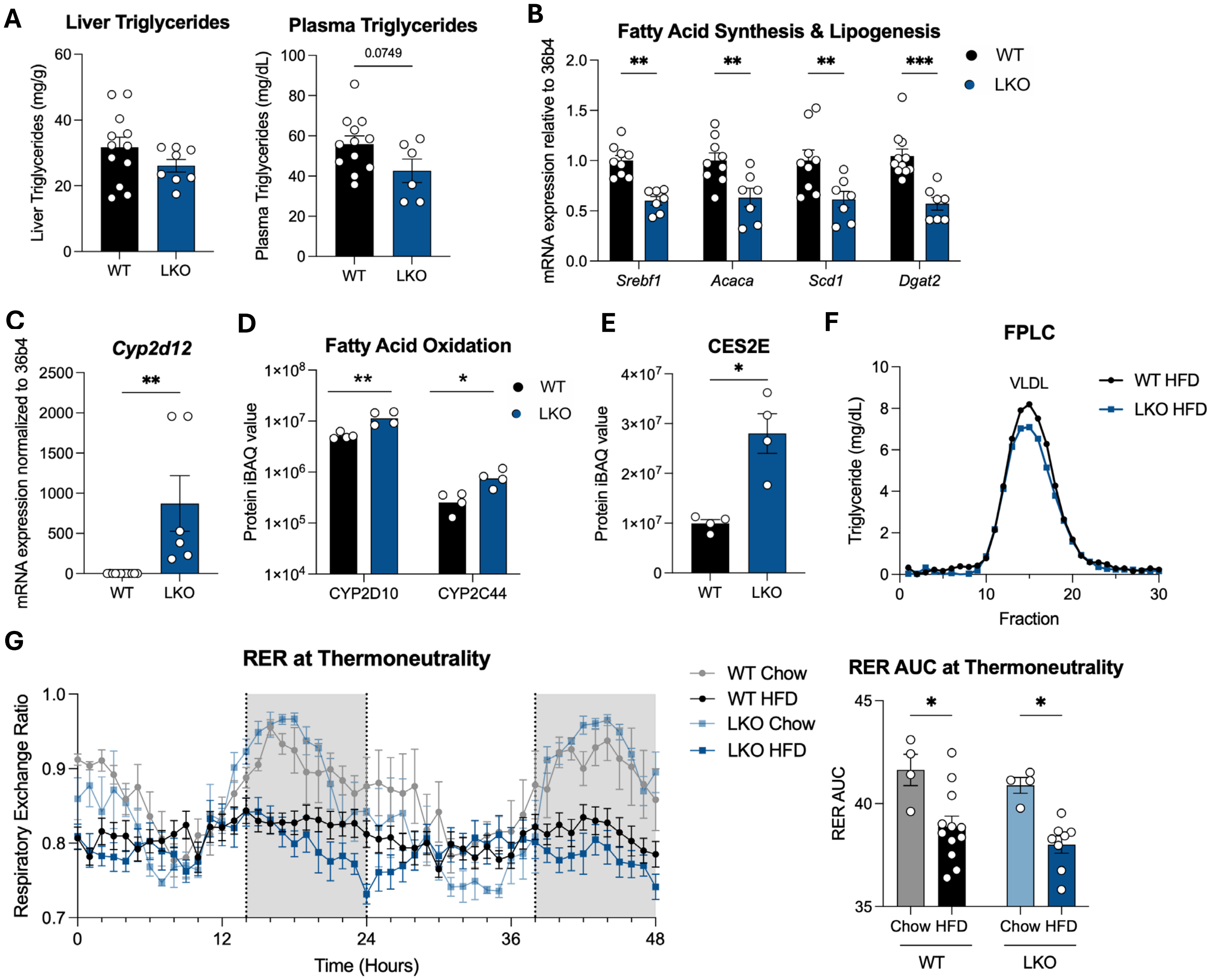


**Figure S9. Decreased expression of genes involved in lipogenesis in male LKO mice compared to WT controls.** (A) Hepatic and plasma triglyceride content was measured in WT and LKO mice. Analysis of fatty acid synthesis/lipogenic (B) and fatty acid oxidation (C) gene expression in HFD-fed WT and LKO mice. The liver proteome was analyzed for proteins involved in fatty acid oxidation (D), and lipid esterase (E) pathways in HFD-fed WT or LKO mice. (F) FPLC of fractionation of fed male HFD-fed WT and LKO mice mouse plasma, pooled from four mice per genotype and measured for triglyceride content. (G) Male HFD- and chow-fed mice were housed in Promethion Metabolic cages at thermoneutrality (30°C), from which oxygen consumption and carbon dioxide production were measured to analyze respiratory exchange ratio (RER). *$P \leq 0.05$, **$P \leq 0.01$, ***$P \leq 0.001$, ****$P \leq 0.0001$, by unpaired two-tailed T-Test (A-E) or two-way ANOVA followed by Holm-Sidak test (G). Data are represented as mean ± SEM with individual animals noted as dots (N=4-12).

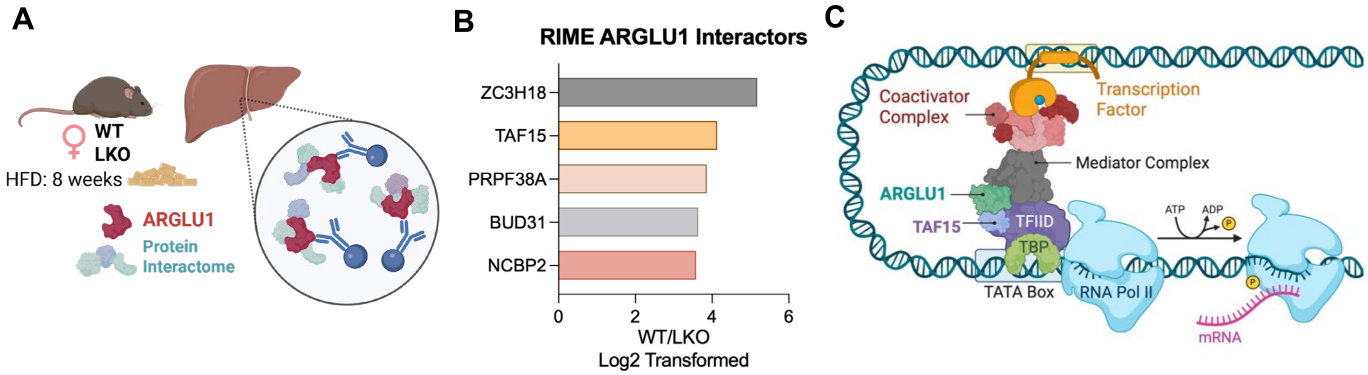


**Figure S10. Protein interactome of ARGLU1 suggesting interaction with splicing factors, transcription factors, and RNA-interacting proteins.** (A) Schematic of Chromatin-Immunoprecipitation Mass Spectrometry experiment design. (B) Top 5 most enriched protein interactors of ARGLU1 from RIME experiment in WT HFD-fed female livers. (C) Proposed mechanism of interaction for transcriptional coregulator ARGLU1.

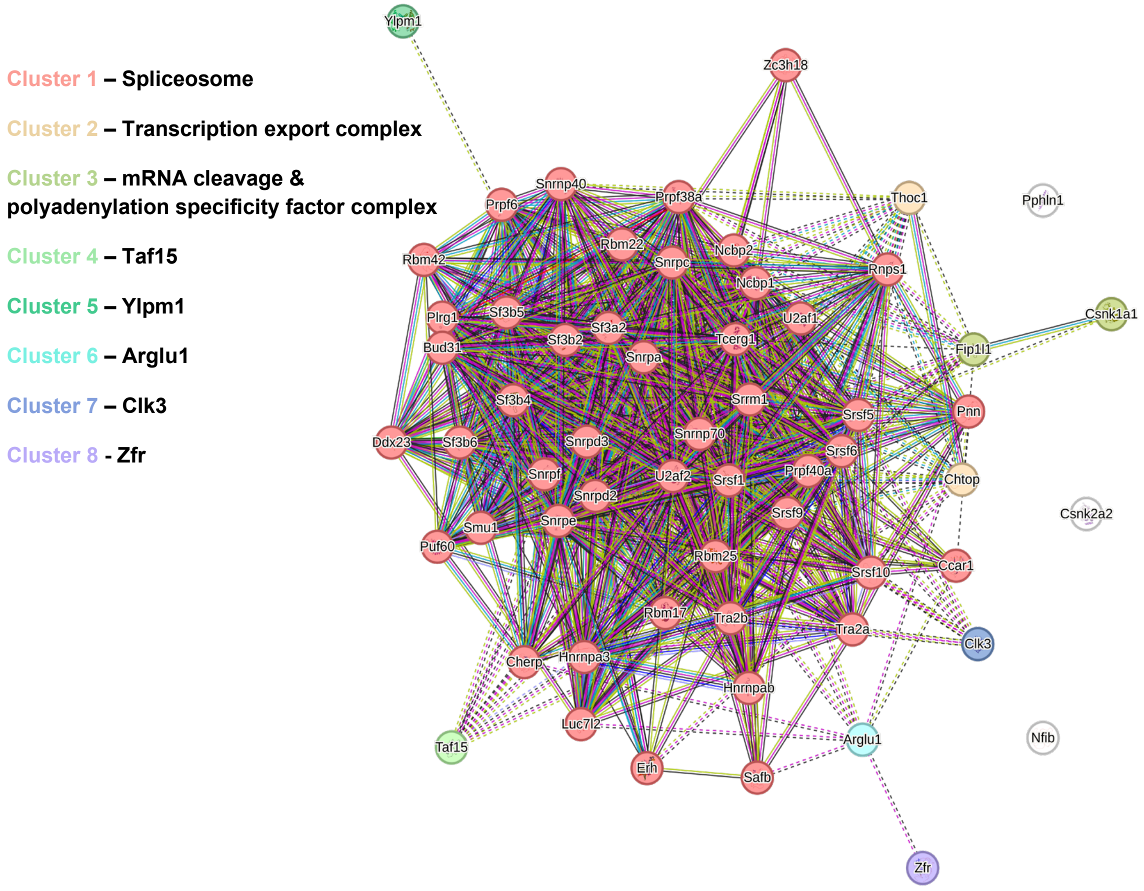


**Figure S11. STRING analysis of 67 enriched ≥1.0 log2-transformed differentially regulated proteins interacting with ARGLU1.** Clustering: K-means clustering.

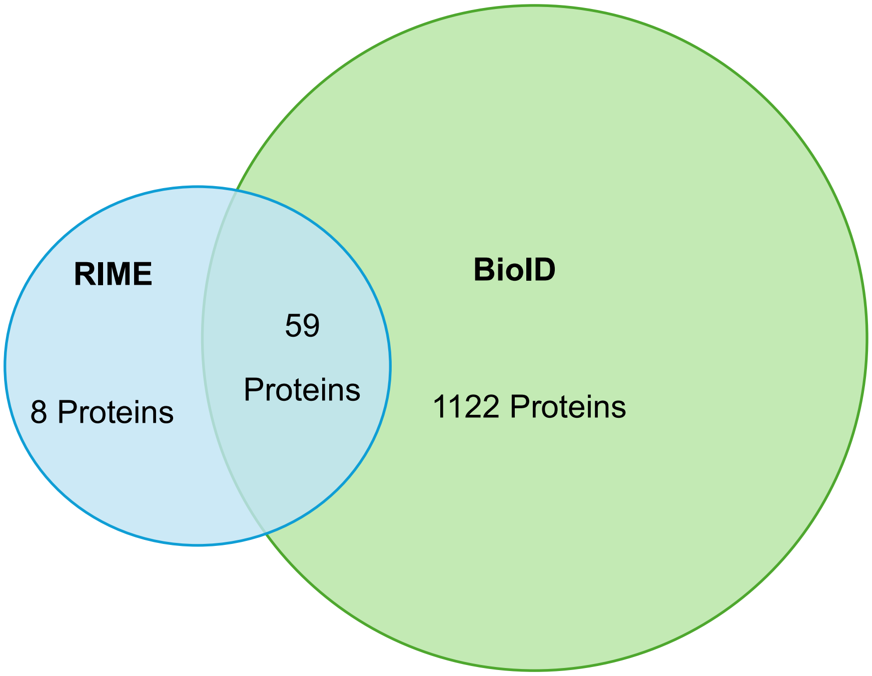


**Figure S12. Venn diagram of proteins identified as interacting with ARGLU1 from RIME and BioID experiments with fold enrichment log2 ≥1.0.** With an enrichment ≥1.0 log2 cut off between ARGLU1-expressing and ARGLU1-null controls, RIME (rapid immunoprecipitation of endogenous proteins using mass spectrometry) identified 67 proteins, while BioID identified 1,181 proteins, with 59 shared between experiments. RIME experiment assessed ARGLU1 binding partners in female HFD-fed WT and LKO liver samples, using an ARGLU1 pull-down. BioID data were collected from HEK293 cells[1]. Overlapped and uniquely regulated genes were found using Venn diagrams (http://bioinfomatics.psb.ugent.be).

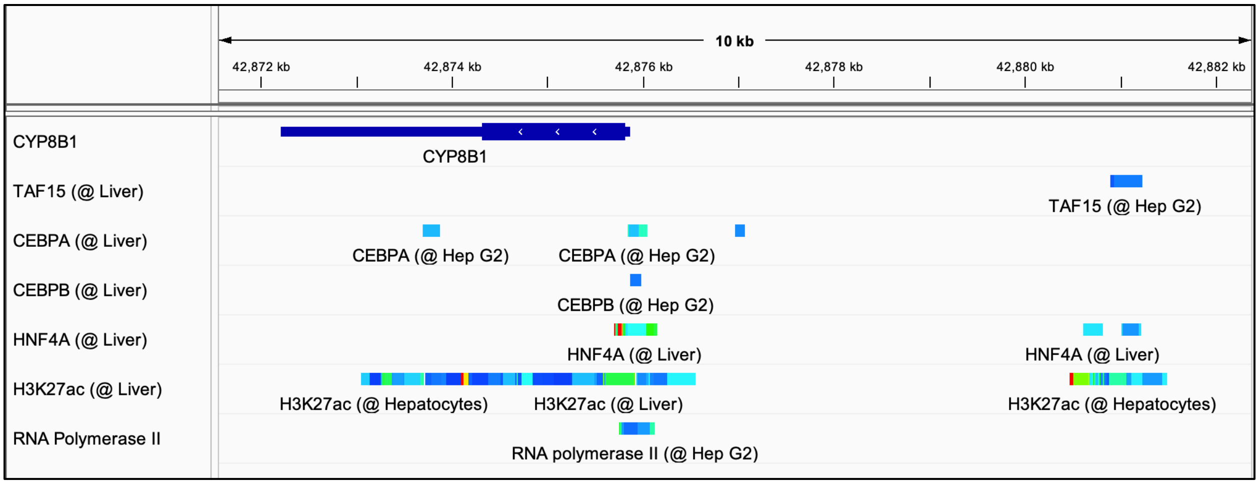


**Figure S13. ChIP-sequencing peaks for C/EBP, HNF4α, and TAF15 in the *CYP8B1* gene from HepG2 cells.** Data collated from ChIP-Atlas and integrated for visualization using Integrative Genome Browser.

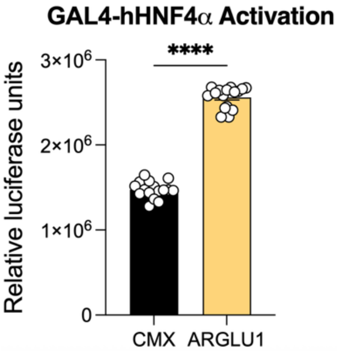


**Figure S14. Increased HNF4α activation (no ligand present) with ARGLU1 co-expression.** HEK293 cells transfected with GAL4-hHNF4α/UAS-luciferase reporter and 15 ng/well of CMX or hARGLU1. **P* ≤ 0.05, ***P* ≤ 0.01, ****P* ≤ 0.001, *****P* ≤ 0.0001, by unpaired two-tailed T-Test. Data are represented as mean ± SEM with individual technical replicates noted as dots (N=16).

**Table S1. Nutrient Information for HFD (TD.88137).**

| | % by weight | % Kcal from |
|---|---|---|
| Protein | 17.3 | 15.2 |
| Carbohydrate<br>(Sucrose) | 48.5<br>(34) | 42.7 |
| Fat | 21.2 | 42.0 |
| Cholesterol | 0.2% | |

**Table S2. Body weight gain (g) on HFD for both WT controls: Albumin $^{Cre/+}$ and Arglu1$^{fl/fl}$, demonstrating similar body weight gain.**

| | Albumin$^{Cre/+}$ | | | Arglu1$^{fl/fl}$ | | | Adjusted *P*-value |
|---|---|---|---|---|---|---|---|
| **Week** | Mean | SEM | N | Mean | SEM | N | |
| **0** | 0 | 0 | 5 | 0 | 0 | 9 | |
| **1** | 1.22 | 0.383 | 5 | 1.044 | 0.416 | 9 | >0.9999 |
| **2** | 1.82 | 0.603 | 5 | 2.366 | 0.466 | 9 | 0.9997 |
| **3** | 3.88 | 0.762 | 5 | 3.633 | 0.583 | 9 | >0.9999 |
| **4** | 5.2 | 0.965 | 5 | 5.333 | 0.727 | 9 | >0.9999 |
| **5** | 7 | 0.985 | 5 | 6.1 | 0.811 | 9 | 0.9904 |
| **6** | 8.06 | 1.02 | 5 | 6.833 | 0.982 | 9 | 0.9381 |
| **7** | 8.76 | 0.952 | 5 | 7.566 | 1.239 | 9 | 0.9467 |

**Table S4. Bile acid standards information.**

| **Abbreviation** | **Full name** | **Vendor** | **CAS** |
|---|---|---|---|
| CA | Cholic acid | CALBIOCHEM | 229101 |
| CDCA | Chenodeoxycholic acid | Sigma | C9377 |
| DCA | Dexoycholic acid | CALBIOCHEM | 264101 |
| HCA | Hyocholic acid | Cayman Chemicals | 20293 |
| LCA | Lithocholic acid sulfate | Sigma | L-5756 |
| HDCA | Hyodeoxycholic acid | Nutritional Biochemical Corporation | U05067 |
| Desoxy-CA | Desoxycholic acid | Miles-Ames Research Division | A503312 |
| UDCA | Ursodeoxycholic acid | Sigma | U-5127 |
| βMCA | Beta-Muricholic acid | Cayman chemical | 2393-5901 |
| ωMCA | Omega-Muricholic acid | Cayman chemical | 6830-03-01 |
| T-CA | Taurocholic acid | Sigma | T-4009 |
| T-DCA | Taurodeoxycholic acid | Sigma | T-0875 |
| T-LCA | Taurolithocholic acid | Sigma | T-7515 |
| T-LCA sulfate | Taurolithocholic acid sulfate | Sigma | T-0512 |
| T-CDCA | Taurochenodeoxycholic acid | Sigma | T-6260 |
| T-αMCA | Tauro- Alpha-Muricholic acid | Cayman chemical | 2260905-08 |

| | | | |
|---|---|---|---|
| T-βMCA | Tauro-beta-Muricholic acid | Cayman chemical | 145022-92-0 |
| T-ωMCA | Tauro-Omega-Muricholic acid | Cayman chemical | 2456348-84-6 |
| T-HDCA | Taurohyodeoxycholic acid | Sigma | 700248p |
| T-HCA | Taurohyocholic acid | Cayman Chemical | 22669 |
| T-UDCA | Taurine Ursodeoxycholic acid | Cayman Chemical | 20277 |
| G-CA | Glycocholic acid | Sigma | G-7132 |
| G-DCA | Glycodeoxycholic acid | Sigma | G-3258 |
| G-CDCA | Glycochenodeoxycholic acid | Sigma | g0759 |
| G-LCA | Glycolithocholic acid | Sigma | G-0134 |
| G-UDCA | Glycoursodeoxycholic acid | Sigma | 6863 |
| G-HDCA | Glycohyodeoxycholic acid | Cayman Chemical | 22643 |
| 7-Keto-DCA | 7-Keto Deoxycholic acid | Cayman chemical | 911-40-0 |
| CDCA-d4 | Chenodeoxycholic acid d4 | Toronto Research Chemicals | C291902 |
| UDCA-d4 | Ursodeoxycholic acid d4 | C/D/N Isotopes | D-3819 |

**Table S6. Primer sequences for qPCR.**

| ***Abbrev.*** | ***Gene Name*** | ***Accession No.*** | ***Sequence (Fwd and Rev, 5'>3')*** |
|---|---|---|---|
| *36b4* | Ribosomal protein, large | NM_007 475.5 | CGTCCTCGTTGGAGTGACA<br>CGGTGCGTCAGGGATTG |
| *Abca1* | ATP-binding cassette subfamily A member 1 | NM_013 454.3 | TCCTCATCCTCGTCATTCAAA<br>GGACTTGGTAGGACGGAACCT |
| *Abcb11* | Bile Salt Export Pump | NM_003742.4 | AAGCTACATCTGCCTTAGACACAGAA<br>CAGAAGAGGTCCACCCTCTC |
| *Abcg5* | ATP Binding Cassette Subfamily G Member 5 | NM_031884.1 | GTCAATGAGTTTTACGGCCTGAA<br>CACATCGGGTGATTTAGCATAGAG |
| *Abcg8* | ATP Binding Cassette Subfamily G Member 8 | NM_026180.2 | GAGCTGCCCGGGATGATA<br>CCCGGAAGTCATTGGAAATCT |
| *Acat2* | Acetyl-CoA Acetyltransferase 2 | NM_009338.3 | CCCGTGGTCATCGTCTCAG<br>GGACAGGGCACCATTGAAGG |
| *Ces1e* | Carboxylesterase 1 | NM_001025194.2 | TGTCTCTGTTCTTGTGTTGT<br>CAGATATGACAGCAATTTTC |
| *Cyp7a1* | Cytochrome P450 Family 7 Subfamily A Member 1 | NM_000780.4 | AAGGAGGACTTCACTCTACA<br>GTCGTATTTAAAAGTCAAAGG |
| *Cyp7b1* | Cytochrome P450 Family 7 Subfamily B Member 1 | NM_001324112.2 | TCTGCCTGGGAATTTAG<br>AAAAGCATAATCAGCAGTAAC |
| *Cyp8b1* | Cytochrome P450 Family 8 Subfamily B Member 1 | NM_004391.3 | CTTACTCCAAATCCTACCAG<br>GAAATTAACAGTCGCACAC |
| *Cyp27a1* | Cytochrome P450 Family 27 Subfamily A Member 1 | NM_000784.4 | CCAGGAACAGGTCAAGA<br>TTGTTCAGCACCTGGA |

| | | | |
|---|---|---|---|
| *Hmgcr* | 3-Hydroxy-3-Methylglutaryl-CoA Reductase | NM_000859.3 | CTTGTGGAATGCCTTGTGATTG<br>AGCCGAAGCAGCACATGAT |
| *Hmgcs1* | 3-Hydroxy-3-Methylglutaryl-CoA Synthase 1 | NM_001098272.3 | GCAGTCTTCAATGCCGTGAA<br>CCTGCAACTACCAGAGCATATCG |
| *Npc1l1* | Niemann-Pick C1-Like 1 | NM_013389.3 | GCAAGGTGATCAGGAGGTTGA<br>ATCCTCATCCTGGGCTTTGC |
| *Slc10a1* | Solute Carrier Family 10 Member 1 (NTCP) | NM_003049.4 | GAAGTCCAAAAGGCCACACTATGT<br>ACAGCCACAGAGAGGGAGAAAG |
| *Soat1* | Sterol O-acyltransferase 1 | NM_001252511.2 | ACTGCAAAGAAGATGAATATC<br>CACATCTGGTTTACCTTTCA |
